# MitoPerturb-Seq identifies common and gene-specific single-cell responses to mitochondrial DNA depletion and heteroplasmy

**DOI:** 10.1101/2025.07.08.663208

**Authors:** Stephen P. Burr, Kathryn Auckland, Angelos Glynos, Abhilesh Dhawanjewar, Wei Wei, Cameron Ryall, Antony Hynes-Allen, Malwina Prater, Matylda Sczaniecka-Clift, Julien Prudent, Patrick F. Chinnery, Jelle van den Ameele

## Abstract

Mitochondria contain their own genome, the mitochondrial DNA (mtDNA), which is under strict control of the cell nucleus. mtDNA occurs in many copies in each cell, and mutations often only affect a proportion of them, giving rise to heteroplasmy. mtDNA copy number and heteroplasmy level together shape the cell- and tissue-specific impact of mtDNA mutations, ultimately giving rise to rare mitochondrial and common neurodegenerative diseases. However, little is known about how copy number and heteroplasmy interact within single cells, and how this is regulated by the nuclear genes and pathways that sense and control them. Here we describe MitoPerturb-Seq for CRISPR/Cas9-based high-throughput single-cell interrogation of the impact of nuclear gene perturbation on mtDNA copy number and heteroplasmy. We screened a panel of nuclear mtDNA maintenance genes in cells with heteroplasmic mtDNA mutations. This revealed both common and perturbation-specific aspects of the integrated stress-response to mtDNA depletion, that were only partially mediated by Atf4, and caused cell-cycle stage-independent slowing of cell proliferation. MitoPerturb-Seq thus provides novel experimental insight into disease-relevant mito-nuclear interactions, ultimately informing development of novel therapies targeting cell- and tissue-specific vulnerabilities to mitochondrial dysfunction.

## Introduction

Mitochondria store their own genetic information in a ∼16.5 kb small circular genome, the mitochondrial DNA (mtDNA), which encodes 37 genes required for mitochondrial oxidative phosphorylation (OXPHOS). Each cell may contain hundreds to thousands of copies of mtDNA, and their replication and transcription are strictly controlled by nuclear-encoded mitochondrial localized proteins. mtDNA copy number (CN), the absolute amount of mtDNA molecules per cell, shows considerable tissue- and cell-type specific variability,^1,2^ and age-dependent decrease in mtDNA CN may contribute to aging and neurodegeneration.^2,3^ Mutations in the mtDNA often only affect a proportion of mtDNA molecules within a cell, a state called heteroplasmy.^4,5^ During development and aging, the number and proportion of mutated mtDNA molecules can increase, to high levels in single cells. Symptoms arise when deleterious heteroplasmy levels reach a cell type-specific threshold, leading to biochemical OXPHOS defects, cell dysfunction and cell death. Age-related clonal expansion of mtDNA mutations has been implicated in the pathogenesis of rare mitochondrial diseases,^6^ and common neurodegenerative disorders, including Alzheimer’s and Parkinson’s disease,^7,8^ as well as in specific types of cancer.^9,10^ Identification of factors and pathways that regulate mtDNA CN and reduce heteroplasmy, even by a few percent below the threshold, offers the possibility to reverse biochemical defects and confer protection against a range of rare and common diseases.

Genome wide association studies have shown the importance of nuclear loci modulating mtDNA CN and heteroplasmy, but little is known about the specific genes involved. Most studies to date have been correlative and were based on bulk tissue analysis containing diverse cell types in varying proportions, with inconsistent and contradictory findings.^4,11–23^ As a result, for many nuclear-encoded genes and genomic loci, a direct causal link between gene activity and mtDNA dynamics remains to be established. Moreover, given the extraordinary degree of mtDNA mosaicism across tissues, with virtually all cells containing different mixtures and levels of wild-type (WT) and mutant mtDNA,^24–28^ it remains unclear how mtDNA CN and heteroplasmy levels interact within individual cells to cause a downstream tissue-specific phenotype.

Here, we describe and deploy MitoPerturb-Seq, allowing us to determine the impact of specific nuclear genetic perturbations on mtDNA CN, heteroplasmy and the resulting transcriptomic response at single-cell resolution. Targeting a library of candidate nuclear genetic modifiers of mtDNA dynamics in heteroplasmic mouse embryonic fibroblasts (MEFs), we show gene-specific modulation of mtDNA CN and heteroplasmy variance, affecting cell-cycle progression and the associated nuclear transcriptional response. DamID-seq-based chromatin profiling confirmed Atf4 as a key mediating transcription factor for some, but not all nuclear genes responding to decreased mtDNA levels, with different ‘mitochondrial stress’ transcriptional signatures linked to specific nuclear genes despite a similar effect on mtDNA CN. MitoPerturb-Seq thus offers a unique opportunity for unbiased high-throughput interrogation of disease-relevant interactions between nuclear and mitochondrial genomes, within single, isogenic cells. This approach controls for multiple confounding factors enabling the delineation of a direct causal link between specific nuclear genes, critical mtDNA parameters and downstream transcriptional responses.

## Results

### Single-cell CRISPR screening with whole-cell Multiome in heteroplasmic cells

We established MitoPerturb-Seq to understand whether disruption of candidate nuclear genes modulates mtDNA at the single-cell level. MitoPerturb-Seq combines CROP-seq for pooled single-cell CRISPR screening^29^ with a 10X Genomics Multiome-based approach^30^ for combined scATAC- and scRNA-seq in whole cells. This enables the simultaneous profiling of mtDNA sequence, CN and heteroplasmy levels from scATAC-seq data, and the detection of gRNA sequences from the single-cell transcriptome (**Fig. 1A;S1A**). We generated Cas9-expressing mouse embryonic fibroblasts (MEFs) (**Fig. S1B**) from heteroplasmic mice carrying an m.5024C>T point mutation in the mitochondrial tRNA^Ala^ gene (*mt-Ta*),^16,31^ which corresponds to the human mitochondrial disease-causing heteroplasmic m.5650G>A tRNA^Ala^ mutation^31–33^ (**Fig. 1A**). Selected Cas9-transgenic MEFs had a mean heteroplasmy of 60.8 ± 5.8 % (mean ± SD), just below the threshold for biochemical defects in most tissues,^11^ with low intercellular variability (**Fig. S1C**). MEFs were transduced with a pooled gRNA library (60 gRNAs, 3 gRNAs per gene, 6 control genes, 3 non-targeting (NT) gRNAs) to perturb 13 nuclear genes previously suggested to affect mtDNA CN and/or heteroplasmy, either through their role in mtDNA maintenance (Akap1^34^, Nnt^18^, Polg^17,35^, Tfam^12,36,37^), mitochondrial membrane remodeling (Dnm1l/Drp1^38^, Mtfp1^22^, Opa1^39–41^, Snx9^42,43^), or mitochondrial biogenesis and mitophagy (Atg5^44^, Oma1^18^, Pink1^34,45^, Ppargc1a^46^, Prkn^45,47^) (**Fig. 1B;S1D;Table S1**). Ten days after transduction, gRNA-expressing cells were processed for whole-cell Multiome (**Fig. 1A;S1E,F**). Following quality control (QC) of raw sequencing reads, barcode processing and cell filtering steps, 5,718 high-confidence single cells were identified, with an average of 5,967 genes and 11,783 unique ATAC-seq peaks per cell; data were visualized by weighted-nearest neighbor (WNN) UMAP after cell-cycle correction (**Fig. 1C-E;S1G-M**). 0.0022 % of RNA-seq reads aligned to gRNAs (**Fig. S2A**) and 15.8 % of ATAC-seq reads to mtDNA, corresponding to 31.7 x mean mtDNA sequencing depth per cell.

**Figure 1.**
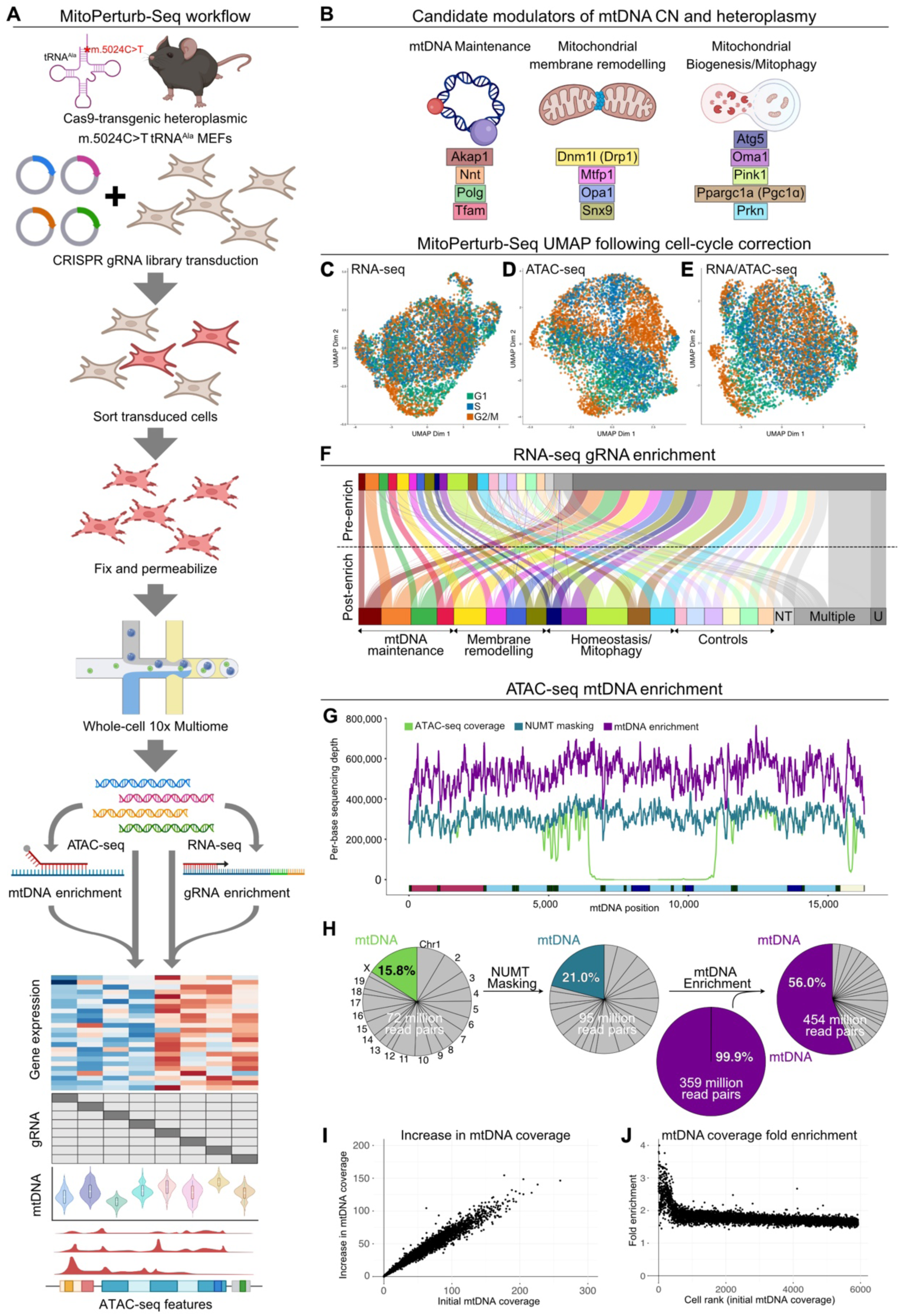
MitoPerturb-Seq combines single-cell CRISPR screening with whole-cell Multiome in heteroplasmic cells. (A) Overview of the MitoPerturb-Seq experimental workflow in heteroplasmic mouse embryonic fibroblasts (MEFs). (B) Candidate genes included in the MitoPerturb-Seq pooled gRNA library. (C-E) Uniform Manifold Projections showing MitoPerturb-Seq cells, clustered based on overall RNA-seq expression (C), chromatin accessibility (D) and a combined Weighted Nearest Neighbor (WNN) analysis (E), following regression of cell cycle heterogeneity from the RNA-seq dataset. Cells are colored by cell cycle phase. (F) MitoPerturb-Seq target gene assignments pre- and post-gRNA enrichment. Target gene colors as in (B); only cells with at least 1 gRNA detected post-enrichment are shown. (G,H) Per-base coverage (G), or percentage of ATAC-seq reads (H) aligning to the mtDNA, following alignment to the standard mm10 genome (light-green), or to a NUMT-masked mm10 genome before (dark green) and after (purple) hybridization-capture based mtDNA enrichment. (I,J) Per-cell absolute (I) or fold increase (J) in mtDNA coverage following mtDNA enrichment, compared to initial pre-enrichment coverage (I) or with cells ordered by initial mtDNA coverage (J). See also Figures S1, S2 and S3.

To improve gRNA assignment, we used Targeted Perturb-seq (TAP-seq)^48^ for PCR-based enrichment of gRNA transcripts. This increased the number of cells assigned to a specific gRNA from 609 (10.2%) to 3,586 (60.1%), expanding the number of cells in each target gene group to 179 ± 41 (mean ± SD) (**Fig. 1F**). Since confidence of estimating true heteroplasmy levels increases with higher read depth for a specific allele (Ref^49^ and **Fig. S2B,C**), we also sought to further increase single-cell mtDNA coverage. We first masked nuclear-localized mitochondrial sequences (NUMTs) in the reference genome to prevent loss of ATAC-seq reads caused by dual alignment to mtDNA and NUMT sequences.^24^ This increased average sequencing depth per cell to 47 ± 37 x (mean ± SD) (21.0% of ATAC-seq reads) (**Fig. 1G,H;S2D,E**). In addition, we performed hybridization capture-based enrichment of mtDNA from scATAC-seq libraries (**Fig. 1A;Table S2**), generating a separate sequencing library consisting of 99.9% mtDNA reads (**Fig. 1H**), with much higher sequencing saturation (mtDNA read duplication rate increased from 38% to 77%) (**Fig. S2F**), evenly spread across the mitochondrial genome (**Fig. 1G;S2G**). This further increased average mtDNA sequencing depth per cell ∼1.7-fold, regardless of initial mtDNA coverage, except in those cells with very low initial coverage (<2x), where depth rose 2- to 3-fold (**Fig. 1I,J**). Together, this resulted in an 80x ± 61x (mean ± SD) average mtDNA sequencing depth per cell (**Fig. S2D**), sufficient to allow reliable mtDNA variant identification and relative CN estimation in most cells (**Fig. S2B,C**). A replicate experiment under identical conditions resulted in 6,272 cells of which 2,965 (43%) could be assigned a unique gRNA identity, with 31.7x ± 30x (mean ± SD) mtDNA coverage after enrichment demonstrating reproducibility of our MitoPerturb-Seq approach in heteroplasmic MEFs (**Fig. S3**).

### MitoPerturb-seq identifies mtDNA depletion following targeted gene perturbation

We combined both experiments into a single integrated dataset (**Fig. S4A,B**) to maximize the potential for discovery of perturbation-related mitochondrial phenotypes. This yielded 11,990 cells with 60.6x average mtDNA sequencing depth. The data were filtered to only retain cells with a unique gRNA assignment, resulting in a total of 6,551 cells. As expected,^50^ target gene knock-down (KD) efficiency for each gRNA was proportional to baseline expression level (**Fig. 2A,B;S4C**). We next assessed how mtDNA CN and heteroplasmy level were affected by the genetic perturbation. Relative mtDNA CN was measured as average single-cell mtDNA coverage from ATAC-seq reads.^2^ Per-cell heteroplasmy level was quantified as proportion of all reads matching either the WT or the mutant mtDNA molecule. The precision of per-cell heteroplasmy measurements was enhanced by combining reads across 7 heteroplasmic single-nucleotide variants (SNVs) (**Fig. S4D-F**), previously shown to be in *cis* or *trans* with the pathogenic m.5024C>T allele,^51^ and confirmed by calculating per-cell correlation of heteroplasmy levels (**Fig. S4G**) and long-read sequencing (**Fig. S4H,I**). When we visualized mtDNA coverage (**Fig. 2C**) and heteroplasmy levels (**Fig. 2D**) on a clustered UMAP plot (**Fig. 2E**), we noted that one cluster (cluster 4) had markedly lower per-cell mtDNA content than the rest of the dataset. This cluster had a distinct gene expression and accessibility profile compared to the other clusters, and was enriched for gRNAs targeting Tfam, Opa1 and Polg (**Fig. 2F-I**). As expected,^35,36,39^ measuring mtDNA CN in each perturbation group confirmed that KD of Tfam, Opa1 and Polg caused a reduction in mean per-cell mtDNA content (**Fig. 2J;S5A**) and a corresponding decrease in mtDNA transcript levels (**Fig. S5B**). Each of the 3 gRNAs targeting these genes caused comparable mtDNA CN reduction, apart from one of the Polg gRNAs (Polg-6) that only slightly decreased Polg expression-levels (**Fig. S4C**; fold change=0.97, p=0.86) and thus had no impact on mtDNA CN, which was removed from downstream analyses. None of the perturbations affected mean single-cell heteroplasmy levels (**Fig. 2K;S5C-E**), possibly reflecting limited power to detect changes of <∼5% (**Fig. S5F**). In keeping with this, mtDNA CN was not different between high vs low heteroplasmy cells, but significantly lower than in cells with mid-range heteroplasmy (**Fig. 2L**).

**Figure 2.**
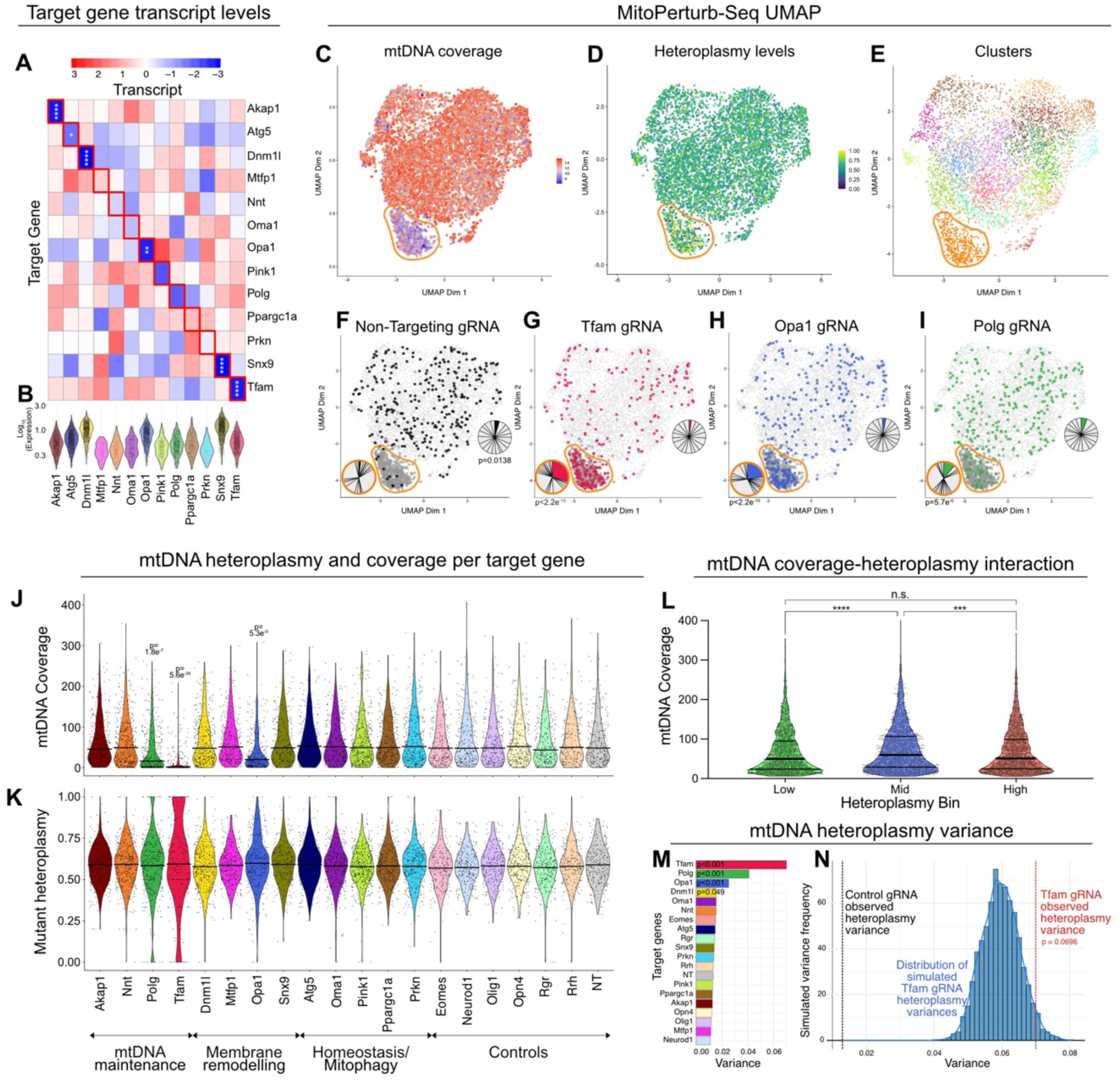
MitoPerturb-seq identifies mtDNA depletion following targeted gene perturbation. (A,B) Scaled transcript levels (A) and per-cell log-transformed transcript counts (B) of candidate target gene across all perturbation groups. Expression values normalized on a per-column basis (A); cells with zero counts are not plotted in (B). Two-tailed t-tests target transcript expression in the relevant perturbation group vs NT gRNA group, * p < 0.05, ** p < 0.01, **** p < 0.0001. (C-E) UMAP shaded according to mtDNA coverage (C), heteroplasmy level (D) and following guided clustering (E). Cluster 4 is highlighted by the yellow outline. (F-I) UMAP with cells assigned to non-targeting (F, black), Tfam (G, red), Opa1 (H, blue) and Polg (I, green) gRNAs highlighted. Pie charts indicate distribution of perturbation groups either in cluster 4 (yellow outline) or all other clusters. Chi-squared tests, p-values in figure. (J,K) Per-cell mtDNA coverage (J) and heteroplasmy (K) for each perturbation group, including controls. Each violin represents the combined data from all three gRNAs targeting the corresponding gene. Pairwise t-tests with multiple testing correction (Bonferroni), p-values specified in figure, all p-values < 0.05 are shown. Horizontal lines indicate median (J) or mean (K). (L) Per-cell mtDNA coverage in low-, mid- and high-heteroplasmy cells. Cells with coverage >50 were ordered by heteroplasmy and split into three bins with equal cell numbers. Median, 1st and 3rd quartiles are shown. Kruskal-Wallis test, **** p <0.0001, *** p < 0.001, n.s. = not significant (M) Variance in heteroplasmy levels for each perturbation group. Brown-Forsythe tests for each group against control gRNAs, with Bonferroni multiple testing correction. (N) Heteroplasmy variance distribution for simulations under a random segregation bottleneck upon Tfam KD, using control gRNA heteroplasmies and Tfam gRNA mtDNA CNs. Empirical two-tailed p-value tests if the observed Tfam variance is consistent with expectations from mtDNA CN reduction alone. See also Figures S4 and S5.

Although there was no difference in the mean value, the heteroplasmy variance was greater in Tfam, Opa1 and Polg KD cells compared to control groups (**Fig. 2M**). *In silico* simulation showed this was consistent with a genetic bottleneck due to the reduced mtDNA CN leading to higher stochasticity in observed heteroplasmy values and an acceleration of random genetic drift, in the absence of selection (p=0.070; **Fig. 2N;S5G**). In addition to technically validating MitoPerturb-Seq, our data confirm that mtDNA CN can be manipulated to modulate the range of heteroplasmy levels within single cells, potentially influencing the number of cells above or below the threshold required to cause an OXPHOS biochemical defect.

### mtDNA depletion affects nuclear gene expression

A key feature of MitoPerturb-seq is the ability to study transcriptional responses to mtDNA depletion at the single-cell level in an isogenic nuclear background. *A priori*, the mechanisms linking Polg, Tfam and Opa1 KD with mtDNA depletion are likely to differ. Polg is primarily involved in mtDNA replication,^52^ Tfam in mtDNA transcription and compaction,^53^ and Opa1 is thought to have an indirect effect on mtDNA maintenance through regulation of mitochondrial membrane dynamics.^39^ We therefore asked whether these pleiotropic effects had an impact on the downstream transcriptional response to mtDNA CN depletion. We identified 203 Tfam-, 118 Opa1-, and 13 Polg-differentially expressed genes (DEGs, log_2_ fold change >0.25; adjusted p-value <0.05), of which 107/215 (50%) were shared between at least two of the three groups and 12/215 (5.5%, all mtDNA-encoded) between all three groups (**Fig. S6A; Table S3**). Interestingly, while mtDNA CN depletion (**Fig. 2J;S6B**) and the reduction in mtDNA gene expression (**Fig. 3A**) caused by Tfam KD were more severe than upon Opa1 or Polg KD, the nuclear gene expression profile of Opa1 KD cells was more similar to Tfam KD than to Polg KD cells (**Fig. 3A,B;S6B**). The differences in nuclear transcriptomic response (nuclear PC1) were more prominent when we controlled for absolute mtDNA CN at the single-cell level (**Fig. 3C**), indicating that, at least for Opa1 KD cells, mtDNA depletion itself was only partially responsible for the nuclear transcriptomic response.

**Figure 3.**
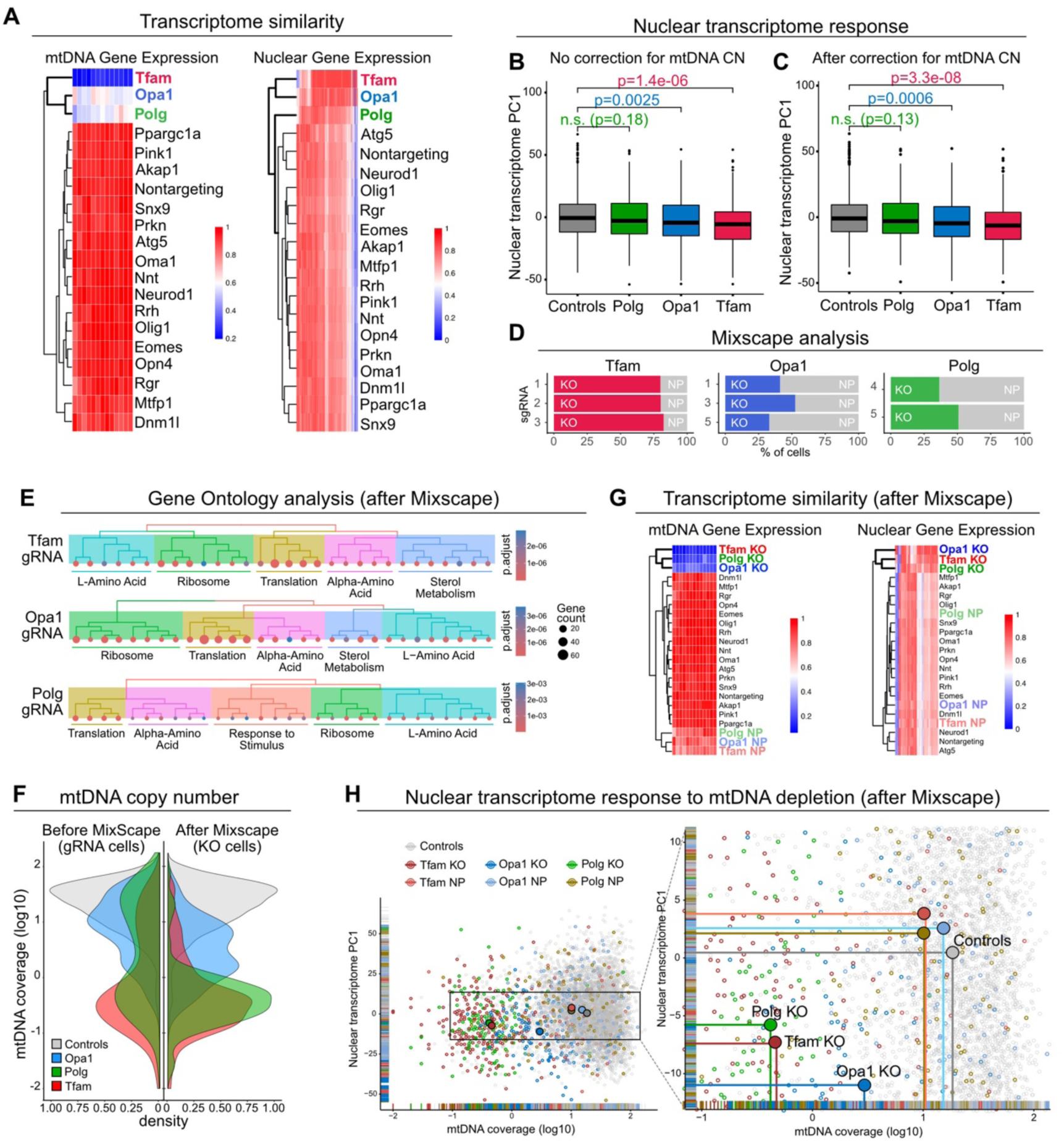
mtDNA depletion affects nuclear gene expression. (A) Average gene expression of selected mtDNA (left) and nuclear (right) genes. Genes with zero expression were removed, filtered for adjusted p-value < 0.05 and scaled to the gene with highest expression. (B,C) Distribution of first principal component (PC1) of nuclear transcriptome across cells in each indicated perturbation group before (B) and after (C) correction for per-cell mtDNA coverage. Wilcoxon signed-rank test, n.s. = not significant. (D) Percentage of cells assigned one of the three gRNAs targeting Tfam (red), Opa1 (blue) and Polg (green) designated as knock-out (KO) following Mixscape analysis. Remaining cells (shaded grey) were assigned as non-perturbed (NP, grey) (E) Significantly enriched Gene Ontology (GO) terms, grouped by biological process, identified in the Tfam, Opa1 and Polg KO groups. Full GO terms analysis is in Table S5. (F) Distribution of mtDNA coverage (log10 transformed) in control cells and the Tfam, Opa1 and Polg perturbation groups before and after removal of NP cells by Mixscape analysis. (G) Average gene expression of selected mtDNA (left) and nuclear (right) genes, following Mixscape classification. As in (A), genes were filtered for adjusted p-value < 0.05 and normalized to the highest-expressing gene. (H) Strength of nuclear PC1 in all cells plotted against mtDNA coverage (log10 transformed), with Polg, Tfam and Opa1 perturbation groups highlighted, separated by KO and NP cells after Mixscape. See also Figures S6 and S7

In order to focus our analysis on the most severely perturbed cells, we used Mixscape^54^ to define the difference between severely perturbed (knock-out, ‘KO’) and less-/non-perturbed (‘NP’) cells within each gRNA-group in an unbiased way (**Fig. 3D;S6C,D**). Mixscape retained more Tfam-KO cells (219/270, 81%) than Opa1-KO (124/308, 40%) or Polg-KO (118/286, 41%) cells, allowing identification of 250 DEGs in Tfam KO cells, 261 in Opa1 KO cells, and 46 in Polg KO cells. 42/325 (13%) DEGs were shared between all three KO groups, and 190/325 (58%) DEGs between at least two of the three KO groups (**Fig. S6E; Table S4**). Gene ontology (GO) analysis of DEGs showed a similar response in all three perturbation groups with an upregulation of mitochondrial integrated stress response (mtISR) pathways, including genes involved in cytoplasmic protein translation, ribosome biogenesis and amino acid metabolism (**Fig. 3E; Table S5**). Sterol metabolism genes were selectively depleted only in Tfam- and Opa1-, but not Polg-KO cells (**Fig. 3E**). Enrichment of genes involved in cholesterol homeostasis has been reported in Tfam KO mouse alveolar macrophages,^55^ further confirming the physiological relevance of our MEF-based screening approach. However, interferon-response genes, previously found to be activated upon Polg or Tfam deficiency^56–58^ or in heteroplasmic mice^59^ were not differentially expressed (**Fig. S6F**). In the remaining perturbation groups, Mixscape analysis only detected ‘KO’ cells in the Atg5 gRNA group, indicating that perturbation of most target genes could not elicit a severe nuclear transcriptomic response in our conditions (**Fig. S6D**). Atg5 KO mainly affected pathways related to lysosome activity, and cytoplasmic protein translation and stability (**Fig. S6G**), with all differentially expressed genes and pathways provided in **Tables S4;S5**.

Interestingly, in the Polg- and Tfam-gRNA groups, Mixscape mostly retained cells (KO) with strong mtDNA depletion, which was not the case for the Opa1 KO group (**Fig. 3F**). This rendered the strongly-perturbed Polg KO cells more similar to Tfam KO than to Opa1 KO cells (**Fig. 3G,H**). This is in keeping with the nuclear response to strong Polg- or Tfam-perturbation being primarily caused by mtDNA depletion, while other factors likely contribute to the response in Opa1 KD/KO cells. Focusing our analysis on mtDNA-encoded gene expression (mtRNA), Opa1 KD caused a more severe decrease in mtRNA for the same level of mtDNA depletion than Polg KD (**Fig. S6H-K**), supporting a role for Opa1 in regulation of mtRNA transcription or stability, independent of its effect on mtDNA content, as previously suggested.^60^

We independently validated these findings using bulk RNA-seq analysis and ddPCR-based mtDNA CN measurements after single gRNA transduction. Tfam KD resulted in a rapid and severe mtDNA CN reduction three days after transduction (**Fig. S7A**), implicating active mtDNA degradation, and in keeping with Tfam playing a crucial role in nucleoid compaction and protecting mtDNA from lysosomal or mitochondrial nuclease-mediated degradation.^61–64^ Nevertheless, the transcriptomic response to Opa1 KD 6 days after gRNA transduction was more severe than to Tfam KD (respectively 82 and 33 DEGs at FDR < 0.05, 27 genes overlap) (**Fig. S7B,C**). Together, these data indicate that, although Opa1-, Polg- and Tfam-perturbation all activate a similar set of mtISR genes, retrograde mito-nuclear signaling may occur through different perturbation-specific pathways, with cells responding to Opa1 KD independent of mtDNA depletion, in line with Opa1’s upstream role in mitochondrial homeostasis and membrane dynamics.^65^ Critically, simultaneously probing these mechanisms in nuclear isogenic cells, grown together and exposed to the same environment, allowed us to avoid additional and potentially unknown confounders impacting cellular stress responses.

### Atf4 only partially contributes to the response to mtDNA depletion

To define the factors driving the transcriptional response to mtDNA depletion in Tfam, Opa1 and Polg KD cells, we performed single-cell regulatory network inference and clustering (SCENIC) analysis.^66^ SCENIC allows inference of gene regulatory networks (regulons) that show differential activity in each of our perturbation groups (**Fig. S8A; Table S6**). We identified 34 regulons that were differentially active in all three KD groups (**Fig. 4A**), including regulons driven by the transcription factors Atf4, Ddit3, Cebpg or Jund, previously reported to be involved in the cellular response to stress in cell culture,^67^ and in Tfam or Opa1 KO mice.^68,69^ Comparing our dataset to a recent systematic review of putative Atf4 target genes,^67^ 24/40 (60%) high-confidence Atf4-responsive protein-coding genes were also differentially expressed in at least one of our Tfam, Opa1 and Polg KO groups (**Fig. S8B**). To independently validate these findings, we used DamID-seq ^70,71^ for genome-wide profiling of Atf4 chromatin binding sites in our heteroplasmic MEFs (**Fig. 4B;S8C**). This identified 5,789 Atf4-binding peaks near 4,477 genes across the genome (**Table S7;S8**). The most highly enriched motifs under Atf4 peaks match the known ATF/JUN/FOS consensus binding sites^72^ (**Fig. S8D**), validating DamID-seq peaks as genuine Atf4-binding sites. Of the 325 DEGs in Tfam/Polg/Opa1 KO cells, 126 (38.8%) were near Atf4-binding sites in heteroplasmic MEFs, with binding mostly occurring at the transcription start sites (TSS) (**Fig. 4C-E;S8E**). However, at our current thresholds, the majority of DEGs upon Opa1 KO (158/261, 60.5%), Tfam KO (159/250, 63.6%) or Polg KO (35/46, 76.1%) were not Atf4-bound (**Fig. 4F,G**), indicating an important, but not exclusive role for Atf4 in responding to mtDNA-related stress. Critically, a range of other non-Atf4 transcription factor regulons were found upon SCENIC analysis (**Fig. 4A**). These are provided for further exploration in **Table S6**, and are likely to co-regulate the response to perturbation of these mtDNA maintenance genes or to mtDNA depletion in heteroplasmic cells.

**Figure 4.**
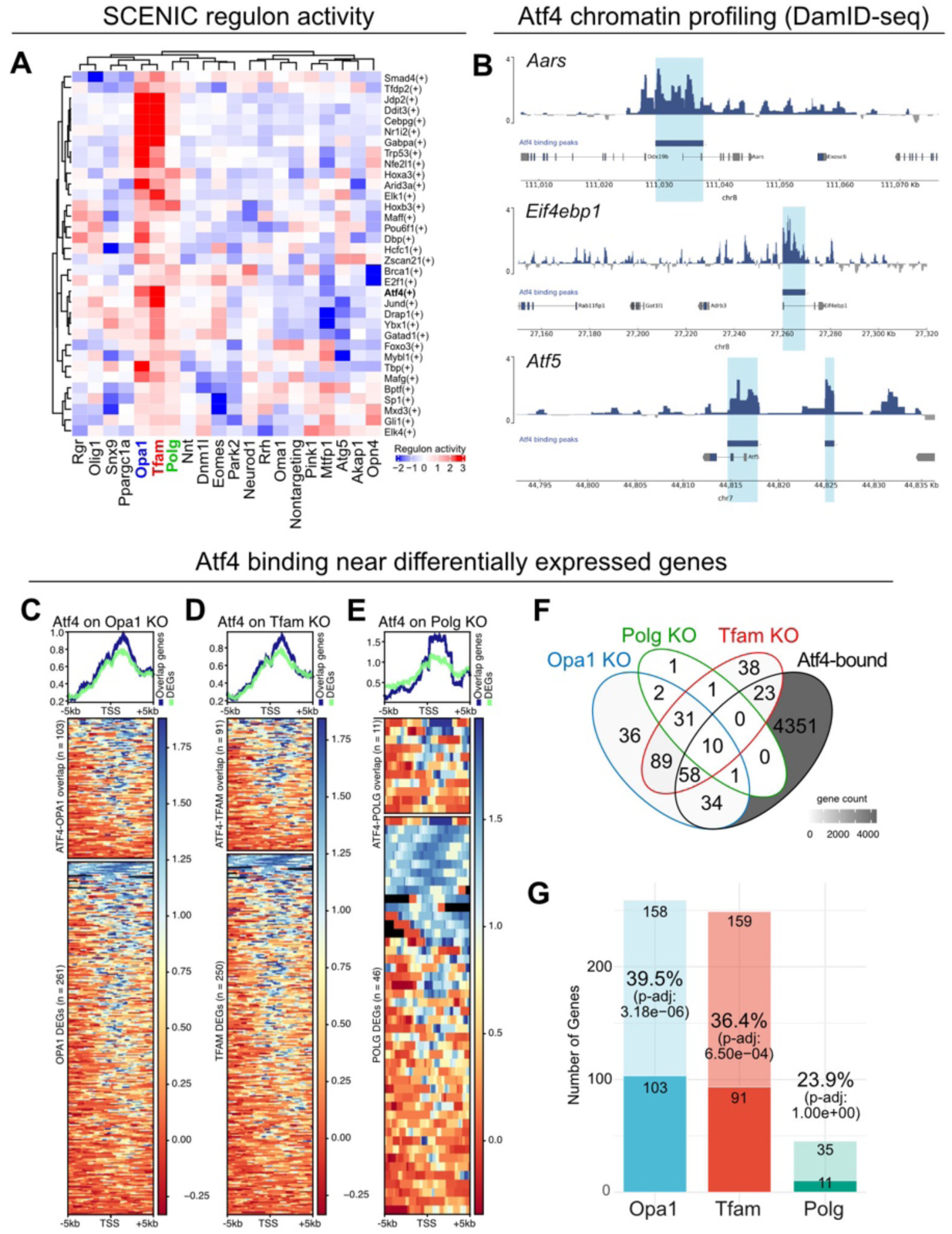
Atf4 only partially contributes to the response to mtDNA depletion. (A) Regulon activity (AUC > 0) for three selected perturbation groups (Opa1, Polg, Tfam) following SCENIC analysis. (B) Atf4 DamID-seq profiles in heteroplasmic MEFs across three Atf4 target genes (*Aars*, *Eif4ebp1* and *Atf5*); Atf4 binding peaks shaded light-blue. (C-E) Metagene plots (top) and heatmaps (bottom) showing Atf4 DamID-seq signal across transcription start sites (TSS) of genes differentially expressed in Opa1 (C), Tfam (D) or Polg (E) perturbation groups, called as associated (top, blue) or not associated (bottom, green) with nearby Atf4-binding sites. (F,G) Overlap between Atf4 DamID-seq target genes and differentially expressed genes in each of the indicated perturbation groups. Fisher’s Exact test with Bonferroni correction, compared to all unique genes detected in gRNA-containing cells. See also Figure S8.

### mtDNA depletion delays cell-cycle progression across all stages

Having identified the transcriptomic changes caused by mtDNA depletion, we next asked how this could impact cellular physiology and behavior. Cell proliferation is a sensitive indicator of mitochondrial activity, with OXPHOS dysfunction previously shown to cause cell-cycle slowing primarily at the G1/S transition.^73–75^ We performed cell-cycle stage annotation on our MitoPerturb-Seq dataset, using Seurat^76^ (**Fig. 5A**) and continuous cell-cycle pseudotime analysis^77^ (**Fig. S9A**). UMAP-analysis of heteroplasmic MEFs prior to cell-cycle correction showed cells mainly segregating based on cell-cycle stage (**Fig. 5A**). To our surprise, we saw no significant differences in the proportion of cells in each cell-cycle phase between the perturbation groups (**Fig. 5B**). After Mixscape analysis, only Opa1, but not Tfam or Polg KO cells, showed a slight (p = 0.034; p.adj = 0.10) increase in the proportion of cells in G1-phase compared to control cells (**Fig. S9B**). This indicates that, in contrast to previous reports describing cell cycle slowing specifically at the G1/S-transition upon OXPHOS dysfunction,^73–75^ there was no selective delay in G1-S progression in Tfam/Polg KO cells, despite severe mtDNA depletion.

**Figure 5.**
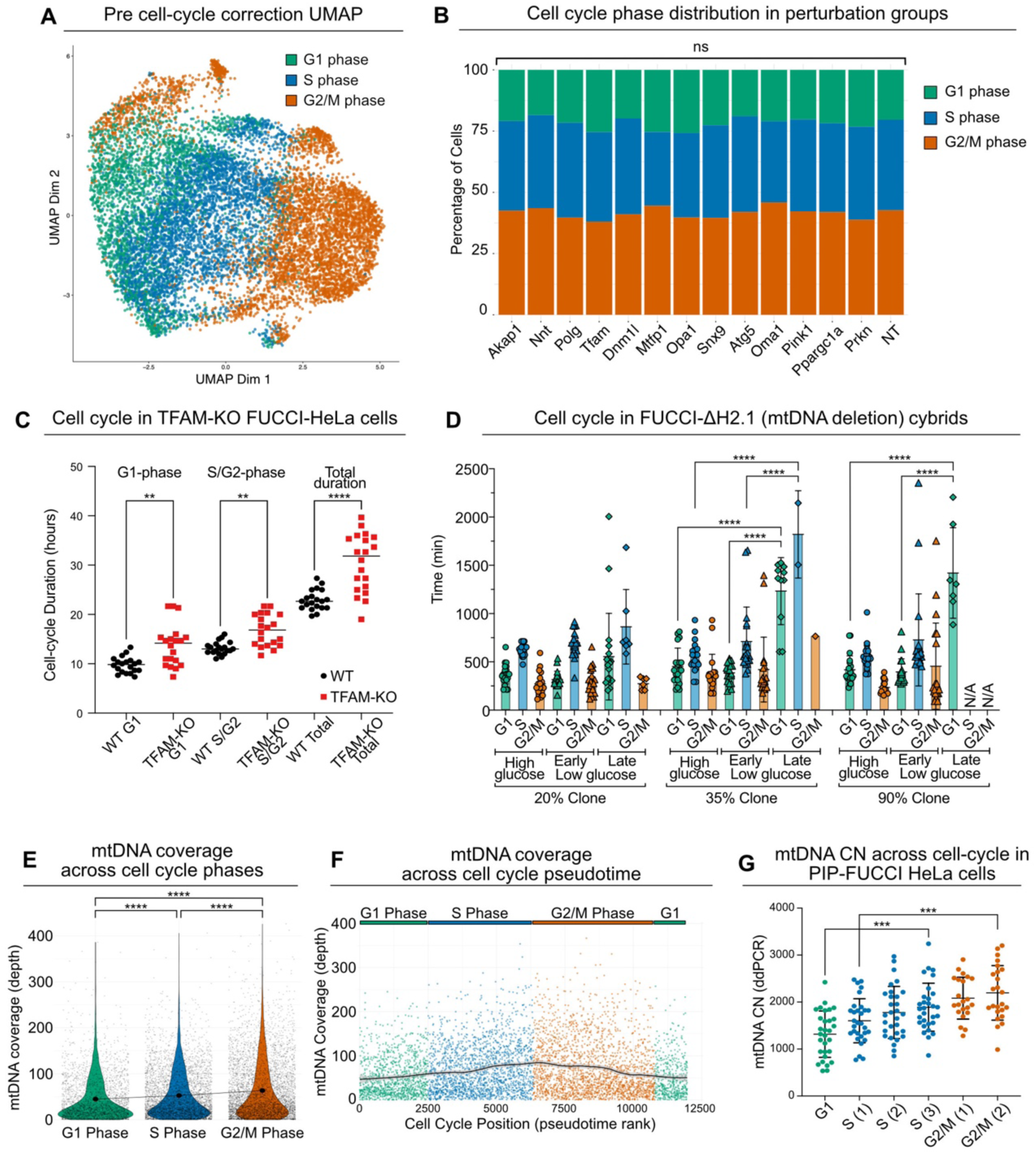
mtDNA depletion delays cell-cycle progression and relaxed replication. (A) UMAP clustering based on RNA-seq prior to regression of cell cycle heterogeneity. Cells are colored by cell cycle phase. (B) Proportion of cells in each perturbation group assigned to each cell cycle stage. Chi-squared tests with multiple testing correction (Bonferroni); ns, not significant. (C) Cell cycle phase duration times in FUCCI-expressing WT and TFAM KO HeLa cells. Comparisons are two-tailed unpaired t-tests, ** p < 0.01, **** p < 0.0001. Error bars are mean ± SD. (D) Cell cycle phase duration in ΔH2.1 mtDNA deletion cybrid clones stably expressing PIPFUCCI, cultured in high or low glucose medium and imaged every 20 minutes over a period of 72h. Phase duration was calculated across one full cycle (G1, S, G2/M) for 20 cells; early, first 24h; late, after 24h. Mean ± SD. One-way ANOVAs per cell cycle phase for each clone with Tukey’s post-hoc test, **** p < 0.0001, all significant pairwise comparisons are shown, N/A = no observed cells progressed to this phase during the experiment. (E) mtDNA coverage across Seurat cell cycle phases. One-way ANOVA with Tukey’s post-hoc test, **** p < 0.0001 (F) mtDNA coverage across cell cycle pseudotime with cells ranked based on tricycle pi score. (G) mtDNA CN in WT PIP-FUCCI HeLa cells flow-sorted by cell cycle phase (sorting strategy in **Figure S9E**). One-way ANOVA with Tukey’s post-hoc test, *** p < 0.001, error bars are mean ± SD. See also Figures S9 and S10.

To validate these findings, we generated transgenic cell-lines expressing a Fluorescent Ubiquitination-based Cell Cycle Indicator (FUCCI)^78^ (**Fig. S9C-E; Supplementary Video 1**) and analyzed cell-cycle duration in WT and TFAM-KO FUCCI-HeLa cells (**Fig. S9F,G**), and in FUCCI-transgenic cybrid cells carrying a heteroplasmic large-scale mtDNA deletion encompassing the major arc^79^ (FUCCI-DeltaH2.1; **Fig. S9H**). TFAM-KO FUCCI-HeLa cells had severe mtDNA depletion (**Fig. S9G**), and displayed an increase in overall cell-cycle length compared to WT FUCCI-HeLa cells, caused by increased duration of both G1 and S/G2 phases (**Fig. 5C**). These findings in TFAM-KO FUCCI-HeLa cells are consistent with our MitoPerturb-seq data from heteroplasmic Tfam- (and Polg-) KO MEFs, where a consistent delay across all stages of the cell cycle would not change the proportion of cells in each individual phase. Clones of FUCCI-DeltaH2.1 cells had no mtDNA depletion (**Fig. S9I**), but carried a range of heteroplasmies (**Fig. S9J**), with higher heteroplasmy levels severely impacting OXPHOS activity (baseline respiration, ATP-linked respiration and maximum respiratory capacity) (**Fig. S9K**). In contrast to TFAM-KO cells, FUCCI-DeltaH2.1 clones with low (20%), mid (35%) and high (90%) heteroplasmy levels all proliferated normally (**Fig. S9L**). This suggests that the remaining spare OXPHOS capacity from few WT mtDNA molecules present in high-heteroplasmy cells is likely sufficient to support normal cell proliferation in glucose-rich medium. However, the proliferation rates of the mid- and high- (but not low-) heteroplasmy cells decreased when cultured in low-glucose medium (**Fig. S9M**). This was mainly caused by an increase in G1-phase duration, with many mid- and high-heteroplasmy cells not progressing from G1 to S-phase when cultured in low-glucose medium for > 48 hours (**Fig. 5D**). Together, these data indicate differential requirements and sensing of mtDNA abundance vs OXPHOS activity to sustain cell-cycle progression, with the G1-S transition being most sensitive to heteroplasmy and OxPhos deficiency. However, when confronted with severe mtDNA depletion, as in Tfam-KO cells, cells will slow cell-cycle progression across all phases.

### Relaxed replication of mtDNA across the cell cycle

During each cell cycle, the nuclear DNA is replicated only once, in a strictly regulated process in S-phase.^80^ In contrast, the mtDNA is thought to undergo relaxed replication, independent of the cell cycle.^81,82^ Prior studies have found conflicting evidence of whether mtDNA replication rates differ between cell cycle stages, but these either lacked temporal resolution, relied on indirect mtDNA CN measurements, or were based on cell cycle synchronization which directly impacts mitochondrial and cellular metabolism.^83–86^ We reasoned that combined RNA- and mtDNA-profiling from a homogeneous population of proliferating cells, as in our MitoPerturb-Seq dataset, could provide an alternative, unbiased approach to characterize cell-cycle-related mtDNA dynamics at the single-cell level. Analysis of mtDNA scATAC-seq read counts across the cell cycle in MEFs showed a progressive increase in mtDNA coverage between G1-S and between S-G2/M stages (**Fig. 5E**), and across continuous cell-cycle pseudotime^77^ (**Fig. 5F**), in keeping with mtDNA replication being fully ‘relaxed’, independent of the cell nuclear state.^81^ Crucially, this also further validates our approach to use ATAC-seq read coverage as a proxy for mtDNA CN, allowing detection of subtle changes in mtDNA content within single cells. Heteroplasmy levels remained constant between cell-cycle phases (**Fig. S10A**) and over cell-cycle pseudotime (**Fig. S10B**), indicating no active selection for or against m.5024C>T tRNA^Ala^ heteroplasmy. The mtDNA depletion caused by Tfam/Polg/Opa1 KO was also present in all cell cycle stages, and across cell-cycle pseudotime (**Fig. S10C-E**). To validate these findings, we measured absolute mtDNA CN by single-cell ddPCR^87^ in WT FUCCI-HeLa and FUCCI-HEK293T cell-lines (**Fig. S10F,G**). This also revealed a linear increase in mtDNA CN, not only between cell cycle phases (G1-S-G2/M) (**Fig. S10F**), but also between early-, mid-, and late-S-phase and between early- and late-G2-phase (**Fig. 5G;S9E**). Interestingly, culturing WT FUCCI-HeLa cells in galactose medium to force respiration via OXPHOS, induced an increase in mtDNA CN, whilst maintaining a linear increase across cell-cycle stages (**Fig. S10G**), indicating that cells actively adjust mtDNA CN in response to higher dependence on mitochondrial ATP-production. Our findings thus validate previous work at high temporal resolution, demonstrating fully relaxed replication of WT and mutant mtDNA molecules across the cell cycle, but modulated in response to bioenergetic requirements.

## Discussion

Single-cell mtDNA analysis in healthy humans,^24^ or in mice and patients with inherited or age-related mtDNA mutations,^25–28^ has found an extraordinary degree of mosaicism, with virtually all cells across tissues and organisms containing different abundance and mixtures of WT and mutant mtDNA. Age-related clonal expansion of mtDNA mutations has been implicated in the pathogenesis of common neurodegenerative disorders like Alzheimer’s and Parkinson’s disease,^7,8^ in mitochondrial diseases,^6^ and in specific types of cancer.^9,10^ Although the phenotypes caused by these mtDNA mutations are inherently mosaic at the cellular level, most studies to date have relied on bulk, tissue-wide analysis, averaging out the single-cell consequences of mtDNA-related heterogeneity, thus limiting the sensitivity to detect disease-relevant phenotypes and mechanisms. Technical challenges to perturb and measure mtDNA at the single-cell level mean that most studies to date have been correlative, and little is known about which nuclear-encoded proteins respond to or regulate changes in mtDNA CN and mutation load in individual cells or cell-types.

Building on single-cell CRISPR-screening methods^29^ and simultaneous single-cell RNA- and mtDNA-sequencing,^30,88^ we developed MitoPerturb-Seq for high-throughput unbiased forward-genetic screening of factors with a direct causal impact on mtDNA CN and heteroplasmy within individual cells. As proof-of-concept, we screened a panel of 60 gRNAs, targeting 13 nuclear-encoded genes known to be involved in mtDNA dynamics, in Cas9-transgenic MEFs from mice carrying a heteroplasmic mtDNA tRNA^Ala^ mutation corresponding to a human disease mutation.^31,33^ Perturbation of three of these genes, Tfam, Opa1 and Polg, caused strong single-cell mtDNA depletion, with large overlap in the single-cell transcriptomic response to each perturbation, driven by comparable transcription factor regulons. One of these transcription factors, Atf4, has been studied extensively as a central factor mediating the response to mitochondrial dysfunction and cellular stress.^89–91^ However, in our data and conditions, only ∼1/3 DEGs were directly bound by Atf4, which, together with our regulon analysis, implies many other factors to be involved in this response. Here we provide a ranked resource of transcription factors involved in modulating mtDNA CN, including established mtISR genes such as two small Maf transcription factors, Maff and Mafg, known to be involved in tissue-specific responses to oxidative stress.^92^ Cell-type specific expression of stress-responsive transcription factors, together with differential chromatin accessibility^93^ are likely to underlie the striking tissue-specific and context-dependent resilience and vulnerability to mitochondrial dysfunction,^94^ so it will be important to extend our approach to other cell types.

We conducted our experiments in heteroplasmic MEFs, and targeted some of the few nuclear-encoded genes that were previously found to regulate heteroplasmy levels in other contexts.^12,17,18,34–36,38,45,47^ Nevertheless, we could not detect significant changes in heteroplasmy levels in any of the perturbation groups. Although technical considerations may to some extent underlie this lack of effect (low target gene expression in MEFs, cell-to-cell heteroplasmy variability, short duration of perturbation), many of these genes will have context-dependent and cell-type/organism-specific functions. Future experiments including more cells, different cell-types, higher mtDNA read-depth and longer time-frames will be required to confidently reveal or exclude genes and processes that modulate heteroplasmy levels – although our findings indicate that the severe gene KD will have very subtle effects at best.

How nuclear and mtDNA replication rates are coordinated during cell growth and proliferation remains poorly understood.^82–84^ We provide strong evidence for completely relaxed mtDNA replication at the single-cell level, independent of cell-cycle stages, in a range of WT, mtDNA depleted, homoplasmic and heteroplasmic human and mouse cell lines. Interestingly, cell-cycle slowing in response to mtDNA depletion affects the entire cell cycle, equally delaying all phases. This differs from the G1/S-specific delay in response to high heteroplasmy levels or targeted OXPHOS dysfunction that we and others have observed,^73–75^ and might indicate the presence of a specific, yet unknown nuclear sensing mechanism to coordinate cell cycle progression across all cell cycle stages with mtDNA abundance or replication.

Our targeted library provides proof of principle for completely unbiased discovery through genome wide CRISPR screening. In addition, PCR- and hybridization capture-based targeted enrichment approaches allow selective sequencing of gRNAs and mtDNA, thus potentially reducing cost and increasing throughput >5 times. Moreover, MitoPerturb-Seq is uniquely placed to conduct screening in disease-relevant post-mitotic cell-types, including *in vivo*^95^ or in human induced pluripotent stem cell-derived organoids. This is particularly relevant, since mtDNA mutations mostly accumulate in and affect post-mitotic cells.^4,94^ The recent advent of mtDNA-modifying technologies to engineer mtDNA point-mutations^96,97^ and deletions^38,98^ in any cell or model system means that MitoPerturb-Seq can now be employed to compare the response to perturbation in isogenic cells with a range of heteroplasmic and homoplasmic mtDNA defects. We envisage this will soon be combined with mtDNA-based lineage tracing, or more affordable combinatorial indexing approaches like SHARE-seq,^99^ for high-throughput simultaneous detection of gRNAs, mtDNA CN and heteroplasmy in millions of heterogeneous cells from mosaic cultures, tissues and organisms. MitoPerturb-Seq also provides a powerful novel forward genetic screening approach to discover the biological mechanisms driving age-dependent mtDNA CN reduction or clonal expansion of damaging mtDNA mutations, and thereby uncover novel druggable targets to treat both rare and common mtDNA-related and neurodegenerative disorders.

## Acknowledgements

We thank all lab members, H. Ma and Y. Crow for helpful discussions, and R. Horvath and H. Biggs for continuous support and interest; S. Jackson for advice on CRISPR screening; C. Lyons for help with cell-cycle analysis; R. Schulte and G. Grondys-Kotarba from the CIMR Flow Cytometry Facility for assistance with cell sorting; the CRUK Cambridge Institute Genomics Core Facility for library preparation, 10X Genomics Multiome and sequencing services; J.B. Stewart, University of Newcastle, for the m.5024C>T mouse and MEFs; F. Merkle for RFP cDNA; C.T. Moraes, University of Miami, for the ΔH2.1 Cybrid cell line.

JvdA is supported by a Wellcome Clinical Research Career Development Fellowship (219615/Z/19/Z), an Evelyn Trust Medical Research Grant (21-25), and a UKRI BBSRC Responsive Mode Research Grant (BB/X00256X/1). PFC is funded by a Wellcome Collaborative Award (224486/Z/21/Z), the UKRI BBSRC (BB/Y003209/1), the Rosetrees Trust (PGL23/100048), and his research is supported by the NIHR Cambridge Biomedical Research Centre (BRC-1215-20014). The views expressed are those of the author(s) and not necessarily those of the NIHR or the Department of Health and Social Care. PFC and JvdA are further supported by a Wellcome Discovery Award (226653/Z/22/Z), a Medical Research Council (MRC) award (MC_PC_21046) to establish a National Mouse Genetics Network Cluster in Mitochondrial Diseases (MitoCluster), and the LifeArc Centre to Treat Mitochondrial Diseases (LAC-TreatMito) under grant no. 10748. LifeArc is a charity registered in England and Wales under no. 1015243 and in Scotland under no. SC037861. MSC was funded by Cancer Research UK (CRUK) Programme Grant C6/A18796 and Discovery Award DRCPGM\100005. JP, PFC and JvdA acknowledge core funding from the UKRI MRC to the MRC Mitochondrial Biology Unit (MC_UU_00028/5, MC_UU_00028/7 and MC_UU_00028/8).

For the purpose of open access, the authors have applied a Creative Commons Attribution (CC BY) license to any Author Accepted Manuscript version arising from this submission.

## Author contributions

SPB, PFC and JvdA conceived the project. SPB, AG, CR, AHA and JvdA designed and conducted experiments. KA, SPB, AD, WW and MP performed bioinformatic analysis. MSC and JP provided essential reagents. PFC and JvdA supervised the project and obtained funding. SPB, KA, PFC and JvdA wrote the paper, and all authors edited and approved the final manuscript.

## Resource availability

Further information and requests for resources and reagents should be directed to and will be fulfilled by the lead contact, Jelle van den Ameele. All sequencing data have been deposited at GEO and will be publicly available as of the date of publication.

## Declaration of interests

The authors declare no competing interests.

## Materials and Methods

### Animal Models and Husbandry

Mouse husbandry was done in a Home Office-designated facility, according to the UK Home Office guidelines upon approval by the local ethics committee (project licenses P6C97520A and PP8565009). The mouse line used in this study was m.5024C>T (Allele symbol: mt-Ta^m1Jbst^, MGI ID: 5902095), bred on the C57BL/6 background.

### Cell Culture and transgenic cell lines

Unless otherwise stated, all cells in this study were maintained in DMEM-high glucose-pyruvate (Gibco, 41966-029) supplemented with 10% fetal bovine serum (Gibco, 16000-044) and 50 μg/mL uridine (Sigma, U3750) at 37°C and 5% CO_2_ in a humidified incubator.

Primary m.5024C>T MEFs were described previously,^16^ isolated from individual E13.5 embryos and immortalized by transfection with the SV40 large T antigen (pBSSVD2005, a gift from David Ron, Addgene plasmid #21826). Cas9-expressing m.5024C>T MEFs were generated by transducing with lentiCas9-Blast lentivirus (Addgene 52962)^100^. Following transduction, cells were selected with 10 μg/mL blasticidin (Sigma, SBR00022). Clonal populations were isolated and heteroplasmy levels assessed by pyrosequencing. To test Cas9 efficiency, m.5024C>T Cas9-blast clones were first transduced with pLJM1-EGFP lentivirus (Addgene 19319)^101^ to stably express eGFP and then transduced with CROPseq-RFP lentivirus expressing eGFP-targeting gRNAs.^102^ Editing efficiency was assessed by flow cytometry, with mRFP expression (i.e. integration and expression of the gRNA cassette) in absence of eGFP expression (i.e. knockout of the transgene) indicating successful editing.

TFAM KO HeLa cell lines were generated using gRNAs and lentivirus described below. Following transduction, cells were selected for transgene expression with 1 μg/mL puromycin (Sigma, P4512). To isolate clonal TFAM KO populations, puromycin-selected cells were subcloned, and TFAM KO was confirmed by western blot. PIP-FUCCI transgenic cells were generated using plasmids and lentivirus described below. Following transduction, cells were subcloned, selecting cells based on the Cdt1-mVenus G1-phase reporter. Clones were checked after expansion by flow cytometry to identify clones showing expected mVenus and mCherry expression patterns for use in downstream experiments.

H2.1 cybrid cells carrying a heteroplasmic mtDNA deletion were obtained from C. Moraes.^79^

For OXPHOS experiments, in addition to standard high-glucose DMEM, cells were maintained in low-glucose or galactose medium. For low-glucose medium, DMEM 1 g/L glucose (Gibco, 31885-023) was supplemented with 4.5 g/L glucose, 10 % fetal bovine serum and 50 μg/mL uridine. For galactose medium, DMEM no glucose (Gibco, 11966-025) was supplemented with 10 mM D-Galactose (Sigma, G0750), 110 mg/L sodium pyruvate (Gibco, 11360-070), 10% dialyzed fetal bovine serum (Gibco, 26400-044) and 50 μg/mL uridine.^103^

Clonal cell populations were obtained by sorting single cells into individual wells of a flat bottomed 96-well culture plate, containing 200 μL of culture medium, using a BD FACSMelody Cell Sorter as described above. One-week post-sort, wells were checked for the presence of a single colony of cells to reduce the risk of obtaining non-clonal populations. Clones were expanded to 24-well plates after 2 weeks, after which further characterization (e.g. heteroplasmy measurement) was performed to select suitable experimental clones.

### Plasmids and gRNA library construction

We modified CROPseq-guide-Puro (Addgene plasmid #86708) lentiviral plasmid^29^ to embed the gRNA sequences in the 3’ UTR of a poly-adenylated RFP transcript, generating the CROP-seq-RFP lentiviral vector. This enables simultaneous enrichment of RFP-expressing transduced cells by FACS and detection of gRNA sequences via 3’ gene expression profiling. A modified RFP cDNA was a kind gift from F. Merkle, but carried a stretch of 7A’s, leading to spurious annealing of oligo-dT primers within the coding sequence (data not shown). For future experiments, we recommend using the CROPseq-mCherry plasmid that we cloned subsequently. gRNA sequences for CROP-seq (**Table S1**) were designed as described previously,^29^ ordered from Twist Biosciences as a ssDNA oligo pool and PCR amplified prior to cloning. 200 femtomoles of the amplified oligo pool were cloned using the ClonExpress Ultra One Step Cloning Kit (Vazyme, C115) into 10 femtomoles of gel-purified CROPseq-mCherry plasmid digested by BsmBI overnight. The cloning mixture was desalted by dialyzing against nuclease-free water using a 0.05 µm membrane filter (Merck, VMWP02500) for 30 minutes. The desalted cloned plasmids were transformed into Endura electrocompetent cells (Lucigen 60242-1) by electroporation in prechilled 1mm cuvettes at 25 μF, 200 0, 1.5 kV before resuspending in 1 ml recovery medium to recover and then plate, or by heat shock in High Efficiency NEB Stable Competent *E. coli* (NEB C3040H). Bacterial colonies (∼43,000, estimated ∼700x per gRNA) were scraped from plates, plasmid DNA extracted using a Plasmid Midi Kit (QIAGEN, 12143), ran on agarose gel and non-recombined plasmid was extracted using a Monarch DNA gel extraction kit (NEB T1020).

The single gRNA sequence for human TFAM^104^ was ordered as desalted oligos (Sigma) with overhangs for cloning, annealed by heating at 95 °C for 2 minutes and cooled at 1 °C/min to room temperature. The annealed oligo was cloned into the LentiCRISPRv2 backbone as previously reported,^100^ and transformed in 10-beta Competent *E. coli* (NEB, C3019H), and plasmid DNA was extracted using the QIAprep Spin Miniprep Kit (QIAGEN, 27104).

Single gRNA sequences for mouse Tfam, Opa1 and non-targeting were ordered as desalted oligos (Sigma), annealed by heating at 95 °C for 2 minutes and cooled at 1 °C/min to room temperature. Annealed gRNAs were cloned into LentiCRISPRv2-RFP670 (Addgene plasmid # 187646) at a 1:5 vector to insert ratio by Gibson assembly using the ClonExpress Ultra One Step Cloning Kit, transformed in 10-beta Competent *E. coli*, and plasmid DNA was extracted using the QIAprep Spin Miniprep Kit (QIAGEN 27104).

The pCAG-mCherry-P2A-i4Dam and pCAG-mCherry-P2A-i4Dam-ATF4 plasmids were constructed by Gibson assembly into pCAG-IRES-GFP, using mCherry as an upstream open reading frame, and Dam from Addgene plasmid #59217^71^ with a C-terminal Myc-tag. The Atf4 gene was amplified from mouse cDNA and cloned in frame into the pCAG-mCherry-P2ADamID construct 3’ of Dam and the Myc-tag. A P2A ribosome-skipping sequence was placed between mCherry and Dam, resulting in overexpression of Dam and Atf4 (i.e. not ‘targeted’^70^), to override post-transcriptional/translational regulation of Atf4. To allow transient transfection of DamID,^93^ an intron was inserted into Dam by ligating oligos with a modified synthetic intron (‘IVS’) from^105^ into the BamHI restriction site within Dam, between the 3rd and 4th helix of the DNA-binding domain of the Dam methylase^106^, and plasmids transformed in *dam^−^/dcm^−^* competent *E. coli* (NEB C2925H).

### Flow Cytometry

Expression of fluorescent markers was assessed using a BD LSRFortessa Cell Analyzer with appropriate laser and filter/detector settings. Post-acquisition data analysis was performed in FlowJo v10. Single-cell sorting was performed using a BD FACSMelody Cell Sorter according to the manufacturer’s instructions. Briefly, samples were initially gated using doublet discrimination to select live single cells based on forward scatter and side scatter parameters. If cells were being sorted based on expression of a fluorescent marker, additional gating was performed to identify marker-positive cells. Cells in the desired gate were then sorted into tubes (bulk sorts) or plates (single-cell sorts), all sorts were performed in ‘Single Cell’ sort mode.

### Lentiviral Production and Transduction

24 hours prior to transfection, HEK293T LentiX cells (Takara, 632180) were plated at 0.75 x 10^6^ cells per well in 6-well plates. Transfections were performed with cells at 80-90% confluent using TransIT-293 transfection reagent (Mirus, MIR2704) according to manufacturer’s instructions. The envelope plasmid (pMD2.G, Addgene plasmid # 12259), vector plasmid and packaging plasmid (psPAX2, Addgene plasmid # 12260) were added to the transfection mix in a 2:3:4 molar ratio. Supernatant containing lentiviral particles was harvested at 48 h post-transfection and passed through a 40 μm filter to remove cellular debris. Harvested virus was stored at −70 °C prior to use in transductions. Cells to be transduced were plated at 5 x 10^4^ cells per well in 24-well plates, an appropriate volume of thawed and pre-warmed lentiviral supernatant was added and the total volume brought to 500 μL per well with fresh culture medium. 3 hours post-transduction an additional 500 μL of culture medium was added to bring the total volume to 1 mL. Cells were passaged to 6-well plates at 24 hours post transduction. To titrate lentiviral stocks, cells were transduced with increasing volumes of viral supernatant as described above. At 7 days, transduced cells were analyzed to assess expression of the transfer plasmid fluorescent marker, which acted as an indication of the transduction efficiency, this was then used to calculate the volume of viral supernatant required to achieve a given proportion of transduced cells.

### Transient transfection and DamID-seq

All transient transfections were performed using Lipofectamine 2000 (Thermo, 11668027). m.5024C>T MEF cells were plated at 2 x 10^6^ cells per well in 6-well plates in 2 mL of serum free media immediately prior to transfection. Following the manufacturers protocol, 1 μg of each plasmid (pCAG-mCherry-p2A-i4Dam-ATF4 or pCAG-mCherry-p2A-i4Dam) prepared in *dam−/dcm−* competent *E. coli* (NEB C2925H) was added to each well in a 6 well-plate. After 5 hours the media was replaced with DMEM plus FBS. 48hrs post-transfection, cells were harvested, the DNA extracted using a QiaAmp DNA Micro kit (Qiagen 56304), and processed for DamID as described previously.^107^ DamID fragments were prepared for Illumina sequencing according to a modified TruSeq protocol. Sequencing was performed as paired end 50 bp reads by the CRUK Genomics Core Sequencing facility on a NovaSeq 6000.

### Whole-cell 10X Genomics Multiome and sequencing

Cells were prepared for 10X Genomics Multiome ATAC + Gene Expression (GEX) Sequencing following a modified version of the standard 10X Genomics nuclei isolation protocol, similar to what was described previously.^108^ Briefly, 5 x 10^5^ cells were fixed in 0.1% formaldehyde (Thermo) for 5 minutes at room temperature, followed by permeabilization in 0.1% NP40 (Thermo) for 3 mins at 4 °C in the presence of RNase inhibitor (Roche) to prevent mRNA degradation. Fixed and permeabilized cells were resuspended in diluted Nuclei Buffer (10X Genomics), counted and adjusted to a concentration of 3000 cells per μL. Following fixing and permeabilization of cells, subsequent steps were performed at the CRUK Cambridge Institute Genomics Core: Transposition/probe hybridization and ATAC/GEX library preparation were performed using the Chromium Next Gem Single Cell Multiome ATAC and Gene Expression Reagent Kit (10X Genomics, PN-1000283) according to manufacturer’s instructions (protocol ref. CG000810 Rev A). Illumina sequencing was performed by the CRUK Genomics Core Sequencing facility on a NovaSeq 6000, with each run on a single lane of an SP flow cell, returning approximately 650-800 million paired-end reads per lane. Sequencing parameters were set as follows: GEX (Read1 28bp; Read2 90bp; i5 index 10bp; i7 index 10bp); ATAC (Read1 50bp; Read2 50bp; i5 index 16bp; i7 index 8bp).

### gRNA enrichment and sequencing

CROPseq gRNA sequences were enriched from 10X Genomics Multiome gene expression cDNA libraries using TAP-seq^48^ as described previously. Briefly, 15μL of cDNA from the 10X GEM-RT reaction was input to a PCR reaction (TAP PCR 1) containing a CROPseq-RFP specific forward primer binding 54 bp upstream of the 3’ gRNA sequence (CROPouter) and a reverse primer binding the Illumina Truseq Read 1 (Partial Read 1), producing an amplicon approximately 550-600 bp long covering the gRNA sequence and retaining the 3’ 10X cell barcode and UMI sequences. To increase specificity, a second nested PCR (TAP PCR 2) was performed on 10ng TAP PCR1 product using a second CROPseq-RFP specific forward primer carrying the Illumina Truseq Read 2 sequence and binding immediately upstream of the CROPseq-RFP gRNA sequence (CROPinner) and the Partial Read 1 reverse primer. 10 ng of TAP PCR 2 product was input to a third PCR reaction (TAP PCR 3) using a forward primer carrying the Illumina P7 sequencing adapter and a 10 bp index sequence binding the Truseq Read 2 sequence (Illumina P7) and a reverse primer carrying the Illumina P5 sequencing adapter and binding the Truseq Read 1 sequence (Targeted 10X). Following TAP PCR 3, enriched libraries were quantified using the KAPA Library Quantification Kit (Roche). Sequencing on the Illumina MiSeq system was performed using the MiSeq Reagent Nano Kit v2, 300 cycles, according to the manufacturer’s instructions. Following quantification, sequencing libraries were normalized to 4nM, denatured, diluted to a final concentration of 10pM and pooled with 1% 12.5pM denatured PhiX control prior to sequencing. Sequencing parameters were identical to the Multiome GEX libraries described above.

### mtDNA enrichment and sequencing

mtDNA sequences were enriched from Multiome ATAC sequencing libraries by hybridization capture, using a custom xGEN Hybrid Capture Panel (IDT) containing 270 probes targeting the mouse mtDNA sequence (average 1 probe every 60 bp) (**Table S2**). Hybridization capture was performed on 500ng of ATAC sequencing library using the xGen Hybridization and Wash Kit (IDT, 10010351) according to the manufacturer’s instructions, including the optional AMPure XP Bead (Beckman Coulter, A63880) DNA Concentration protocol steps. Post-capture PCR amplification was performed for 12 cycles and the final libraries were quantified using the KAPA Library Quantification Kit and pooled for Illumina sequencing with parameters identical to the Multiome ATAC libraries described above.

### Bulk RNA-seq of single-gRNA CRISPR cells

Following lentiviral transduction with CROPseq-RFP single-gRNA CRISPR vectors, 50,000 RFP-positive cells per sample were sorted at day 6 post-transduction. Total RNA was extracted from the cells using the *Quick*-RNA Microprep Kit (Zymo Research, R1050) and RNA concentration and RIN^e^ values were assessed using the High Sensitivity RNA ScreenTape System (Agilent, 5067-5579/5580). RIN^e^ > 8.5 was confirmed for all samples and samples were adjusted to a final concentration of 10 ng/μL prior to library preparation using the NEBNext Single Cell/Low Input RNA Library Prep Kit (NEB, E6420S) according to manufacturer’s instructions. Libraries were pooled and sequenced on the Illumina NovaSeq X platform on a single lane of a 1.5B flow cell using 50 bp paired-end reads, yielding approximately 750 million reads.

### mtDNA ddPCR and pyrosequencing

Single-cell mtDNA CN measurements were made using the Bio-Rad QX200 AutoDG Droplet Digital PCR system as described previously.^87^ Briefly, single sorted cells in 96-well plates were lysed in lysis buffer containing 1% Tween-20 (Life Technologies, 003005) and 200 μg/mL Proteinase K (Ambion, AM2546) at 37°C for 30 mins followed by 85°C for 15 mins to inactivate Proteinase K. Cell lysate was then input to ddPCR reactions containing primer/probe combinations targeting the mt-Nd1 and mt-Co3 genes of the mouse mtDNA. Following data acquisition, the CNs obtained from the two independent mtDNA probes were averaged to give a final absolute mtDNA CN measurement for each cell.

Single-cell pyrosequencing was performed as previously described,^109^ using the PyroMark Q48 Autoprep pyrosequencing system (QIAGEN) according to the manufacturer’s instructions using a primer set specifically targeting the m.5024C>T mutation site.

### Long-range PCR and long-read sequencing

Genomic DNA was extracted from ear clips using the Monarch Genomic DNA Purification Kit (NEB, T3010) and quantified by Qubit fluorometer. A segment of mtDNA spanning positions 13200 to 5500 was amplified from 10 ng of genomic DNA by long-range PCR using PrimeSTAR GXL Premix (Takara, R051B). The amplicons were isolated by Ampure XP bead cleanup and sequenced by long-read Nanopore sequencing (Plasmidsaurus/Oxford Nanopore Technologies). Reads were aligned to the GRCm39 mouse mtDNA reference genome excluding reads shorter than 8 kb. Variant positions 13614, 13715, 1781, 1866, 3009, 3823, and 5024 were extracted and converted into a binary matrix with the mutant or wild type allele. For each pairwise combination of these positions, the percentage of positions matching either the wildtype or mutant haplotype was calculated to find the co-occurrence of alleles belonging to each haplotype.

### Western Blotting and antibodies

Cultured cells were washed, dissociated using trypsin (Gibco, 15400-054), and pelleted prior to being snap-frozen in liquid nitrogen. Protein extraction was performed by mixing 500μl of PathScan Sandwich ELISA Lysis Buffer (1X) (Cell Signaling, 7081) with each cell pellet. Following a 2 min incubation on ice, samples were spun down for 1min at 14,000 g at 4 °C to remove cell debris. Protein concentration was measured using the Pierce BCA Protein Assay Kit (Thermo, 23227). NuPAGE gels were run at 165 V for 45 min. Membrane transfer was performed using an iBlot machine (Life Technologies). 1hr incubation at RT in 5 % milk in TBS-Tween 0.1 % was used for blocking. The membrane was incubated with the primary antibody overnight at 4 °C, while the secondary incubation was performed at RT for 1 hr. Primary antibodies were anti-TFAM (Cell Signaling, 8076S) and anti-Vinculin (Sigma, V4505). The Clean-Blot IP Detection Kit (HRP) (Thermo, 21232) was used for detection and imaging was conducted using the Amersham Imager 600 (General Electric).

### High resolution respirometry

High resolution respirometry was carried out using the O2k-Respirometer (Oroboros Instruments). Calibration was performed with DMEM plus 10% FBS. 5 million cells suspended in 2 ml of media were added to each chamber. Once the oxygen consumption rate (OCR) reached a plateau, three drugs were added sequentially. First, 5 μl of oligomycin A (Merck, O4876). After establishing the new baseline, 2 μl of carbonyl cyanide m-chlorophenyl hydrazone (CCCP) (Merck, C2759) was added, followed up by successive doses of 1 μl until a plateau of the maximum respiratory capacity was reached. Finally, 1 μl of rotenone (Merck, R8875) and 2 μl of antimycin A (Merck, A8674) were added in quick succession. Baseline respiration was calculated by subtracting the background (respiration still present after the addition of rotenone and antimycin-a from the basal respiration of the sample, prior to the addition of any drugs). Proton-leak was measured by subtracting the background from the respiration still present when oligomycin A was added. ATP-linked respiration was estimated by subtracting the proton-leak from the basal respiration. Finally, the maximal respiratory capacity of the sample was calculated when the background was subtracted from respiration in the presence of CCCP.

### Bioinformatic analysis

#### Initial Analysis and quality control

Raw FASTQ files from Multiome sequencing were combined with the lostread FASTQ files to recover missing or low-quality reads that were initially discarded during sequencing. The FASTQ files were first run through FastQC (v0.11.9, https://www.bioinformatics.babraham.ac.uk/projects/fastqc/) to perform quality control checks on raw sequencing data, and through FastQ Screen (v0.14.1, https://www.bioinformatics.babraham.ac.uk/projects/fastq_screen/) to check for contamination. The combined FASTQ files were then processed using Cell Ranger ARC (v2.0.1) from 10X Genomics, which performs alignment, filtering, barcode counting, and peak calling, to generate feature-barcode matrices for downstream analyses. Cell Ranger ARC was first run with the standard Mus musculus reference genome (GRCm38/mm10, GENCODE vM23/Ensembl 98),^110^ before running with a custom-built Mus musculus reference genome created with `cellranger mkref`. Some mtDNA regions exhibit low coverage due to homology with nuclear DNA and hard-masking these nuclear genome regions (NUMTs) is recommended.^24^ A custom blacklist (https://github.com/caleblareau/mitoblacklist/) was incorporated into the original reference, generating a hard-masked, modified Mus musculus genome. Using the `gex_possorted_bam.bam` file, generated from the Cell Ranger ARC pipeline, the data was run through Qualimap (v2.2.1)^111^ to evaluate the overall mapping quality of our sequencing data.

The filtered feature barcode matrices, stored as sparse matrices in hdf5 format, were loaded into R (v4.3.3) to perform data quality control and standard pre-processing steps with the Seurat R package (v5.1.0),^76^ for RNA data processing, and the Signac R package (v1.13.0),^112^ for ATAC data processing. Data quality control was performed for both scRNA-seq and scATAC-seq data. For RNA, this involved removing cells with; total RNA reads less than 1,000 and greater than 60,000; number of detected genes less than 1,000 and greater than 10,000; mitochondrial RNA percentage greater than 10; and ribosomal percentage greater than 30. For ATAC, this involved removing; total ATAC reads less than 1,000 and greater than 150,000; number of accessible regions less than 500 and greater than 55,000; nucleosome banding pattern signal greater than 2; and transcription start site enrichment score less than 1.

Following quality control, RNA-seq and ATAC-seq data were independently normalized and scaled, after which linear dimensional reduction was performed. For the RNA-seq data, cell cycle scoring was conducted to minimize the impact of cell cycle heterogeneity, assigning each cell a score based on the expression of G2/M and S phase markers. This was initially calculated with the Seurat R package (`CellCycleScoring`), followed by pseudotime analysis with the tricycle R package (v1.12.0)^77^ (described below).

As with quality control, filtering and initial data processing, integration was performed separately for RNA-seq and ATAC-seq data before being combined into one Multiome object for downstream analyses. RNA-seq data integration was performed by first merging the two Seurat objects, followed by `SCTransform` normalization while regressing out cell cycle effects.

Next, linear dimensional reduction was applied, and cell clustering was determined by computing k-nearest neighbors and constructing a shared nearest neighbor graph. Finally, Uniform Manifold Approximation and Projection (UMAP) was used for dimensionality reduction and visualization. Data integration was performed using the `IntegrateLayers` function with the Canonical Correlation Analysis (CCA) method. CCA was selected because the datasets shared common cell types while potentially containing technical or batch-related differences. This method is particularly effective for correcting subtle batch effects while preserving strong biological signals. Following integration, linear dimensional reduction, clustering, and UMAP visualization were repeated. ATAC-seq data integration began by merging the two Seurat objects, followed by linear dimensional reduction, clustering, and UMAP visualization. Integration was then performed using `FindIntegrationAnchors` and `IntegrateEmbeddings`, after which linear dimensional reduction, clustering, and UMAP visualization were repeated. The two integrated Seurat objects were combined, and the `FindMultiModal-Neighbours` function was used to construct a weighted nearest neighbor (WNN) graph, followed by UMAP visualization.

### gRNA detection

gRNA sequences were present in the 90 bp Read 2 sequences of the GEX library, these reads were extracted and assigned to individual cells using a bespoke pipeline: first, raw GEX FASTQ files were processed with UMI-tools (v1.0.1)^113^ `extract` to embed the 10X cell barcode sequence contained in Read 1 into the corresponding Read 2 read name string. Barcoded Read 2 FASTQs were then aligned using bwa (v0.7.17)^114^ `mem` to a custom reference genome containing the sequence for each of the 60 gRNAs in the CROPseq library, flanked on both side by 85 bp anchor sequences from the surrounding CROPseq-RFP backbone, ensuring sufficient reference sequence to successfully align all 90 bp Read 2 sequences that overlapped at least 5 bp of the gRNA sequence. The resulting SAM output was processed in R: aligned reads were filtered to retain those with MAPQ ≥ 30 and individual reads were assigned to single cells by cross-referencing the embedded barcode sequence in the QNAME SAM field with the corresponding barcodes allocated by Cell Ranger ARC. Cells were given a single gRNA assignment if they had ≥ 2 separate barcoded gRNA reads identified and, in cases where multiple gRNAs were identified, the ratio of reads from the most abundant gRNA to total gRNA reads was > 2:3.

### Heteroplasmy calling

Single-cell mtDNA variant identification was performed using the mgatk package (v0.7.0)^24^ implemented in `tenx` mode, with the NUMT-masked Cell Ranger ARC `atac_possorted_bam.bam` output file and corresponding known cell barcode list as inputs, which includes removal of PCR duplicate reads performed at the level of individual cells to give single-cell de-duplicated per-base mtDNA coverage data. High-confidence heteroplasmic variants were reported in the `mgatk.variant_stats.tsv` output. To call single-cell heteroplasmy values, per-base mtDNA sequencing data contained in the `mgatk_signac.rds` output file was first imported into the Seurat object containing the corresponding Cell Ranger ARC GEX and ATAC assays using the `ReadMGATK` Seurat command. Next, base calls at the seven high-confidence heteroplasmic variants corresponding to the reference and mutant alleles were extracted from the mgatk assay and the ratio of mutant to wild-type alleles at each variant position was used to calculate heteroplasmy. Investigation of the correlation between the heteroplasmy calls at individual variant positions, combined with previously published data,^115^ confirmed that the two principal variant positions, m.5024C>T and m.13715C>T, were linked on a single mtDNA haplotype and were inversely correlated with the remaining five variants, m.1781T>C, m.1866G>A, m.3009T>G, m.3823C>T and m.13614T>C, all linked on a second mtDNA haplotype. Based on this confirmed linkage between all seven variants, we were able to treat the coverage at each separate variant position as an independent observation of the same underlying heteroplasmy, thus allowing us to combine the wild-type and mutant base calls at all seven variant sites to increase the effective coverage and maximize the accuracy of the single-cell heteroplasmy calls.

### Heteroplasmy modelling

*In silico* modelling of heteroplasmy calling based on sub-sampling of a simulated heteroplasmic mtDNA population indicated that the sample size, corresponding to the read depth at heteroplasmic sites, has a strong influence on the accuracy of single-cell heteroplasmy estimation, with greater depth resulting in increasingly accurate calls. Previous studies have suggested implementing a minimum coverage cut-off of 20 reads to confidently identify a heteroplasmic variant.^24,116^ To model the effect of mtDNA coverage on the accuracy of heteroplasmy calls, we generated *in silico* cell populations with simulated heteroplasmy values based on the distribution of heteroplasmy in our MitoPerturb-seq dataset. Based on our experimental data we used an mtDNA CN of 1,750, and a mean heteroplasmy of 54% with SD of 12% to generate a set of normally-distributed ‘true’ heteroplasmy values. To investigate the impact of sampling heteroplasmy at different mtDNA coverage, we simulated a population of 2,000 cells and randomly sub-sampled each cell with sample sizes of 5, 20, 50 and 100. These samples were then used to calculate ‘sampled’ heteroplasmy values for the cell and, for each sample size, the sampled heteroplasmy was plotted against the simulated true heteroplasmy for the corresponding cell, with regression line and R^2^ value calculated using ggplot2 (v3.5.1)^117^ `geom_smooth()`. To model the likely impact of applying a coverage threshold of 20 reads to our integrated MitoPerturb-Seq dataset, we simulated a matching population of 6,551 as described above, and sampled these cells using the same distribution of post-mtDNA enrichment coverage that we saw in our integrated data to calculate the ‘sampled’ heteroplasmy value for each cell. Sampled heteroplasmy was then plotted against the simulated true heteroplasmy, with cells sampled at depth < 20 highlighted. Together, this allowed us to conclude that, in our case, a threshold of ≥ 20 reads at the combined heteroplasmic variant sites represented a good compromise, effectively eliminating the cells with the most inaccurate heteroplasmy calls whilst retaining the majority of cells for downstream analysis. We note that in datasets with higher average per-cell mtDNA coverage (e.g. from cells with high mtDNA CN like hepatocytes ^26^), it may well be possible to increase this read-depth threshold to improve overall accuracy without significantly impacting the number of cells retained in the dataset.

### Simulating the effect of reduced sequencing depth on heteroplasmy variance

To test whether the increased heteroplasmy variance observed in Tfam KD cells could be attributed solely to the reduced mtDNA depth at relevant genomic sites, we performed a computational simulation modelling the expected heteroplasmy variance from sampling heteroplasmy levels in control gRNA cells at mtDNA depths characteristic of the Tfam KD cells. For each simulation, cells from the control gRNA group were sampled without replacement to match the number of cells in the Tfam KD group. For each sampled control group cell, a mtDNA depth value was sampled with replacement from the Tfam KD group to approximate the mtDNA depth distribution. A simulated alternative allele count was estimated for each cell via binomial sampling, with the sampled Tfam mtDNA depth as the number of trials and the heteroplasmy level from the sampled control cell as the probability of picking the alternative allele. The heteroplasmy level for each cell was then calculated as the ratio of its simulated alternative allele count to its assigned sampled depth. The null distribution of simulated heteroplasmy variances was generated from 5000 simulations performed as described above and the observed Tfam KD heteroplasmy variance was assessed against this distribution using a two-sided empirical p-value at significance level of α = 0.05.

### Cell-cycle pseudotime analysis

In addition to performing cell-cycle annotation with Seurat, we used the tricycle R package (v1.12.0)^77^ to predict, analyze and visualize cell-cycle states in scRNA-seq. Unlike traditional cell-cycle scoring methods that rely on discrete phase markers, tricycle estimates a continuous cell-cycle trajectory using a reference-based projection approach. By leveraging a pre-trained, internal, reference, using the `Runtricycle` function, tricycle first projects data onto the cell-cycle embeddings using the reference, and then estimates cell-cycle position. The estimated cell cycle positions range from 0 to 2 pi, with 0.5 pi the start of S stage, pi the start of G2M stage and 1.5 pi the middle of M stage. The results of this function are added as metadata to the Seurat object. To evaluate tricycle’s performance, we examined the expression of key genes relative to cell cycle position. Cell cycle annotations from Seurat and tricycle showed high concordance, with 82.3% of annotations aligning. Of the 17.7% that did not align, 9.7% of these were a mismatch between G1 and S annotation, probably because Seurat assigns S and G2M scores based on the expression of predefined marker genes, with cells exhibiting low scores for both phases inferred to be in G1 phase. Additionally, chi-square tests for independence with Bonferroni multiple testing correction were performed for each gRNA across the different cell-cycle phases to assess association between gRNA presence and cell-cycle phase distribution.

### Mixscape analysis

For unbiased perturbation assessment from scRNA-seq datasets, we used the Mixscape method,^54^ implemented within the Seurat R package. We first calculated perturbation signatures (`CalcPerturbSig`), setting the number of nearest neighbors to 20; when clustering by these signatures, technical variation is removed and a specific perturbation cluster is identified. Using these signatures, the `RunMixscape` function assumes each target gene is a mixture of two Gaussian distributions (knock-out (KO) and non-perturbed (NP)), and that NP cells have the same distribution as those expressing non-targeting gRNAs (NT). Each cell is assigned a posterior probability of belonging to the KO group; cells with a probability greater than 0.5 are labelled KO. For the present study, Mixscape analysis was only conducted in cells that were already confidently assigned a gRNA, without cells labelled as either unknown or containing multiple gRNAs during prior gRNA assignment. All cells labelled with a negative control gene (Eomes, Neurod1, Olig1, Opn4, Rgr, Rrh) or non-targeting gRNA were used as the control population. Following Mixscape assignment, the `PlotPerturbScore` function was used to examine posterior probabilities and perturbation score distributions, comparing those assigned as KO or NP to those assigned NT. Differential expression analyses were performed and visualized with the `MixscapeHeatmap` function to see if KO cells exhibited reduced expression. In order to maximize class separability, dimensionality reduction was performed with linear discriminant analysis (LDA) and visualized.

### Differential gene expression

Before differential gene expression, the `PrepSCTFindMarkers` function from the Seurat R package was used to prepare SCT-normalized data for differential gene expression analysis when using the `FindMarkers` set of functions. For the present data, the `FindAllMarkers` differential expression analysis function in Seurat was used to identify marker genes for all clusters in the single-cell dataset. This involved comparing each gRNA group to the control group (negative control genes and non-targeting gRNAs) to identify genes that are significantly expressed in each group. Differential expression analyses were also conducted for cells classified by the Mixscape analysis as KO, comparing each gRNA with a KO classification to the control group (negative control genes and non-targeting gRNA). Differential gene expression results were filtered for an adjusted p-value less than 0.05 and a log2 fold change of 0.25.

Functional enrichment analysis was done using the clusterProfiler R package (v4.12.1).^118,119^ Gene ontology (GO) enrichment analysis was done with the `enrichGO` function, compared to a background gene set (org.Mm.eg.db, v3.20.0). After performing `enrichGO`, the enrichPlot R package `pairwise_termsim` function with the Wang method^120^ was applied to evaluate the semantic similarity between the enriched GO terms based on the topological structure of the GO graph. Relationships between the enriched GO terms were visualized as a hierarchical tree using the `treeplot` function from enrichPlot.

Nuclear principal component analysis (PCA) was conducted using RNA-seq data after integration, normalization and initial filtering. A linear model was fitted to the data, with corrections applied for mtDNA coverage as estimated from the ATAC-seq data. The top 2000 variable residuals from this model were utilized for nuclear PCA. Additionally, nuclear PCA was performed prior to the correction for mtDNA coverage.

### Single-Cell rEgulatory Network Inference and Clustering (SCENIC)

Raw scRNA-seq counts from the QC filtered and integrated dataset were used as an input to the Python (v3.12.2) based pyScenic pipeline (v0.12.1):^121^ Gene regulatory network (GRN) inference was performed using the GRN2Boost algorithm, followed by regulon prediction with the pyscenic `ctx` command with the `–mask_dropouts` set to TRUE. Finally, AUCell (v1.28.0)^66^ was used to calculate cellular regulon enrichment scores, which were then used for downstream analysis. The resulting loom file was loaded into R using SCopeLoomR (v0.13.0, https://github.com/aertslab/SCopeLoomR). The expression matrices, regulons and AUCell matrix were extracted using `get_dgem`, `get_regulons`, and `get_regulons_AUC`, respectively; key column attributes and metadata were also extracted. Cells were split by gRNA and for each group of cells, the AUC matrix was extracted and the mean regulon activity across all of those cells was calculated. The resulting matrix was scaled by performing z-score scaling per regulon and regulons with missing values were removed. To examine the regulon activity scores for select genes, only regulons with a relative activity score greater than zero in all three genes of interest (Opa1, Polg and Tfam) were retained for visualization. Heatmaps were generated with ComplexHeatmap (v2.18.0).^122^

### Bulk RNA-seq analysis

Overall sequencing quality of Raw FASTQ files was first checked using FastQC, confirming no major issues. Next, reads were trimmed to remove Illumina TruSeq adapter sequences using Trimmomatic (v0.39)^123^ and aligned to the mouse GRCm38/mm10 genome reference using RUM (v2.0.4)^124^. Aligned bam files were used as input to HTSeq-count(v2.0.3)^125^, using `union` mode, to count reads in features. Final analysis was performed in R using the EdgeR software package (v4.4.2)^126^.

### DamID-seq analysis

Raw sequencing reads from Dam-only and Dam-Atf4 samples were processed using damid-seq_pipeline (v1.5.3).^127^ Paired-end reads were aligned to the GRCm38/mm10 reference genome using Bowtie2 (v1.3.1)^128^, and mapped reads were assigned to GATC fragments. Signal intensities were computed in 300 bp bins using RPM (reads per million) normalization. Each Dam-Atf4 sample was normalized against each Dam-only sample, resulting in six pairwise comparisons. Peak calling on each pairwise comparison was performed using MACS3 (v3.0.3)^129^ with the Dam-Atf4 sample used as the treatment and the corresponding Dam-only sample used as control. Peaks were called in `broad` mode, with a fixed fragment size of 300 bp and no model estimation (`--nomodel`). The effective genome size was computed from the mm10 reference genome, and significance thresholds were set at p < 0.05 with an mfold range of 5 to 50. Reproducible peaks from the pairwise comparisons were identified across the biological replicates using IDR (Irreproducible Discovery Rate),^130^ broad peaks for each Dam-Atf4 sample were merged across Dam-Atf4 vs Dam-only comparisons and de-duplicated based on signal strength. IDR was computed between all pairwise combinations of the three Dam-Atf4 samples using the python IDR package (v2.0.3) and peaks with a reproducibility score below an IDR threshold of 0.01 were retained. A multi-sample intersection was then performed using bedtools (v2.31.0)^131^ `multiinter` generating a consensus peak set that captured overlapping peaks across replicates. The resulting peaks were annotated using HOMER (v5.1)^132^ `annotatePeaks.pl`, and motif enrichment analysis was performed using HOMER `findMotifsGenome.pl` with a window size of 300 bp around the peaks to identify enriched sequence motifs within the peak regions. Statistical significance of intersections between Atf4 binding targets and Differentially Expressed Genes was performed using the R package SuperExactTest (v1.1.2)^133^ and visualization plots were generated using ggplot2^117^ and ggVennDiagram (v1.5)^134^. GO enrichment analysis was performed on genes associated with Atf4 binding sites using the clusterProfiler R package. The `enrichGO` function was used to identify significantly enriched biological process (BP) terms with a q-value threshold of 0.05. The background gene universe was set to 17,693, the number of genes detected in the 6,551 cells that were assigned a gRNA, and relationships between enriched GO terms were visualized as a hierarchical tree plot using the `treeplot` function in the enrichPlot R package (v1.28.2). Gene Set Enrichment Analysis (GSEA) was conducted using clusterProfiler `gseGO` to assess the ranking of Atf4 target genes within biological pathways. Atf4 binding signal on the differential expressed genes was visualized as heatmaps using the `plotHeatmap` function from deepTools (v3.5.6).^135^ Genomic tracks for Atf4 binding signals from DamID-seq were plotted using pyGenomeTracks (v3.9).^136^

### Quantification and statistical analysis

Unless described otherwise, data visualizations were generated with inbuilt package functions from the Seurat R package, including `DimPlot`, `FeaturePlot` and `VlnPlot`; with `ggplot2`,^117^ alongside `ggplot2` extensions: `ggalluvial` (v0.12.5),^137^ `ggpmisc` (v0.6.1), `ggpubr` (v0.6.0), and `ggvenn` (v0.1.10); and with `ComplexHeatmap`,^122^ `enrichplot`, and `pheatmap` (v1.0.12). Statistical tests were conducted in R; details of the tests can be found in the accompanying figure legends. Figures in this publication have been created with BioRender under license.

## Supplementary Figure legends

**Figure S1.**
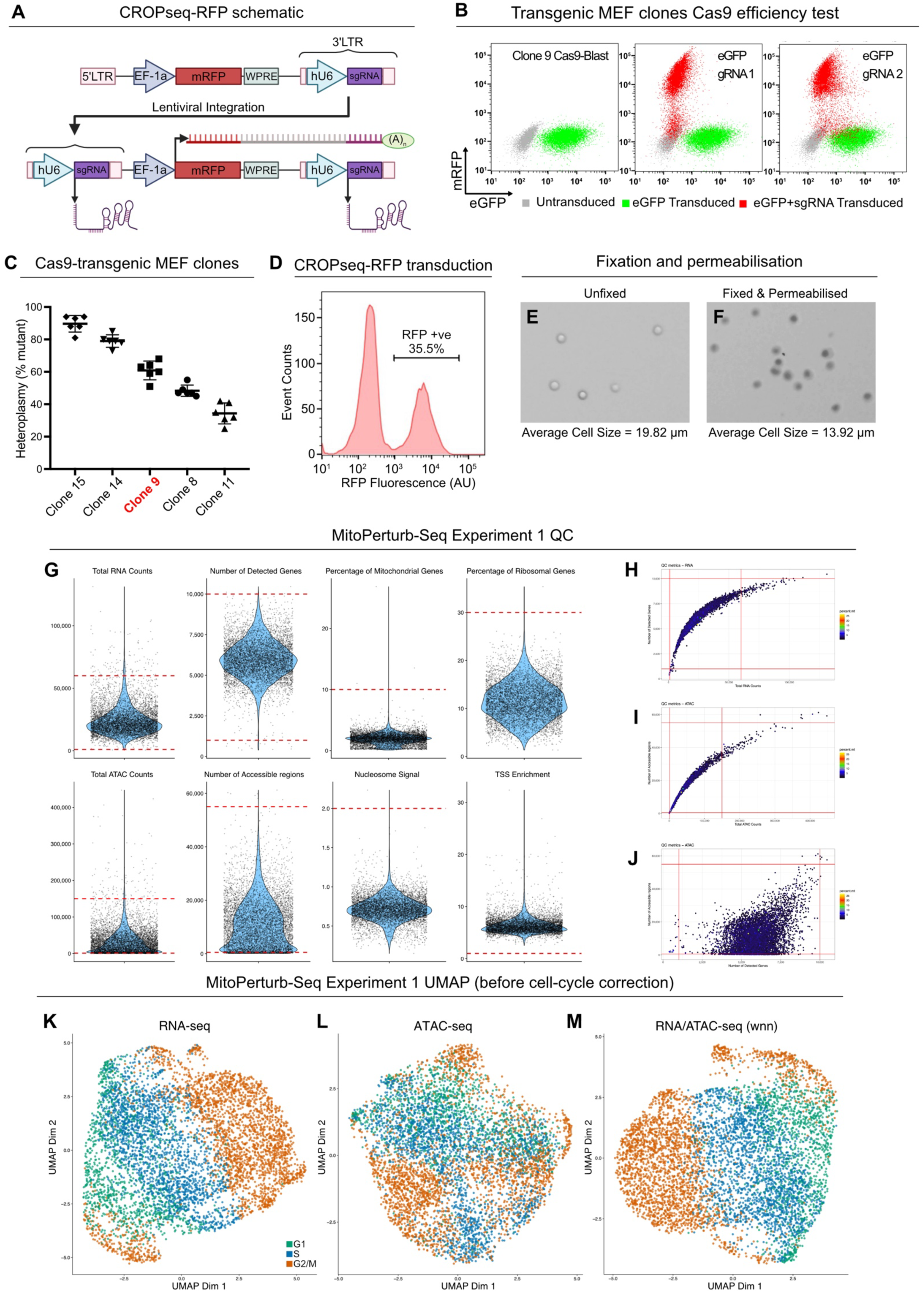
MitoPerturb-Seq and quality-control steps. (A) Schematic of the CROPseq-RFP lentiviral cassette showing transcript expression and polyadenylation of a gRNA-containing transcript following genomic integration. (B) Flow cytometry plots confirming genome editing efficiency in transgenic Cas9-expressing MEF clones. Clonal cells stably expressing eGFP were transduced with CROPseq-RFP lentivirus carrying two separate gRNAs targeting the eGFP sequence. RFP-high cells displayed loss of eGFP fluorescence, confirming effective CRISPR-Cas9 editing of the target gene. (C) Single-cell heteroplasmy measurements from five Cas9 transgenic MEF clones, six cells per clone. Clone 9, used for MitoPerturb-Seq experiments, is highlighted in red. Error bars are mean ± SD. (D) Flow cytometry plot showing RFP expression 10 days post-transduction with the pooled CROPseq-RFP gRNA library. Transduction efficiency of 35.5% (MOI = 0.35) indicates that the majority of cells received a single viral particle. (E,F) Bright field images of unfixed/unpermeabilized (E) or fixed/permeabilized (F) MEF cells stained with trypan blue. Viable cells (E) have not taken up the dye and remain bright with a well-defined membrane, while permeabilized cells (F) have taken up the dye, with darker-staining nuclei. Average cell size, calculated by the Countess cell counter, is indicated. (G) QC violins from Seurat for RNA-seq and ATAC-seq, dashed red lines indicate thresholds used for filtering, as described in the methods. (H-J) QC plots of nCountRNA (H), nCountATAC (I) and nFeatureRNA (J) versus nFeatureRNA (H) or nFeatureATAC (I,J), colored by percent mitochondrial reads, with lines indicating cutoff thresholds shown in violin plots in (G). (K-M) UMAPs showing MitoPerturb-Seq cells, clustered based on overall RNA-seq (K), ATAC-seq (L) or by Weighted Nearest Neighbor (WNN) of combined RNA- and ATAC-seq (M) data, prior to regression of cell cycle heterogeneity. Each point represents an individual cell. Cells are colored by cell cycle phase.

**Figure S2.**
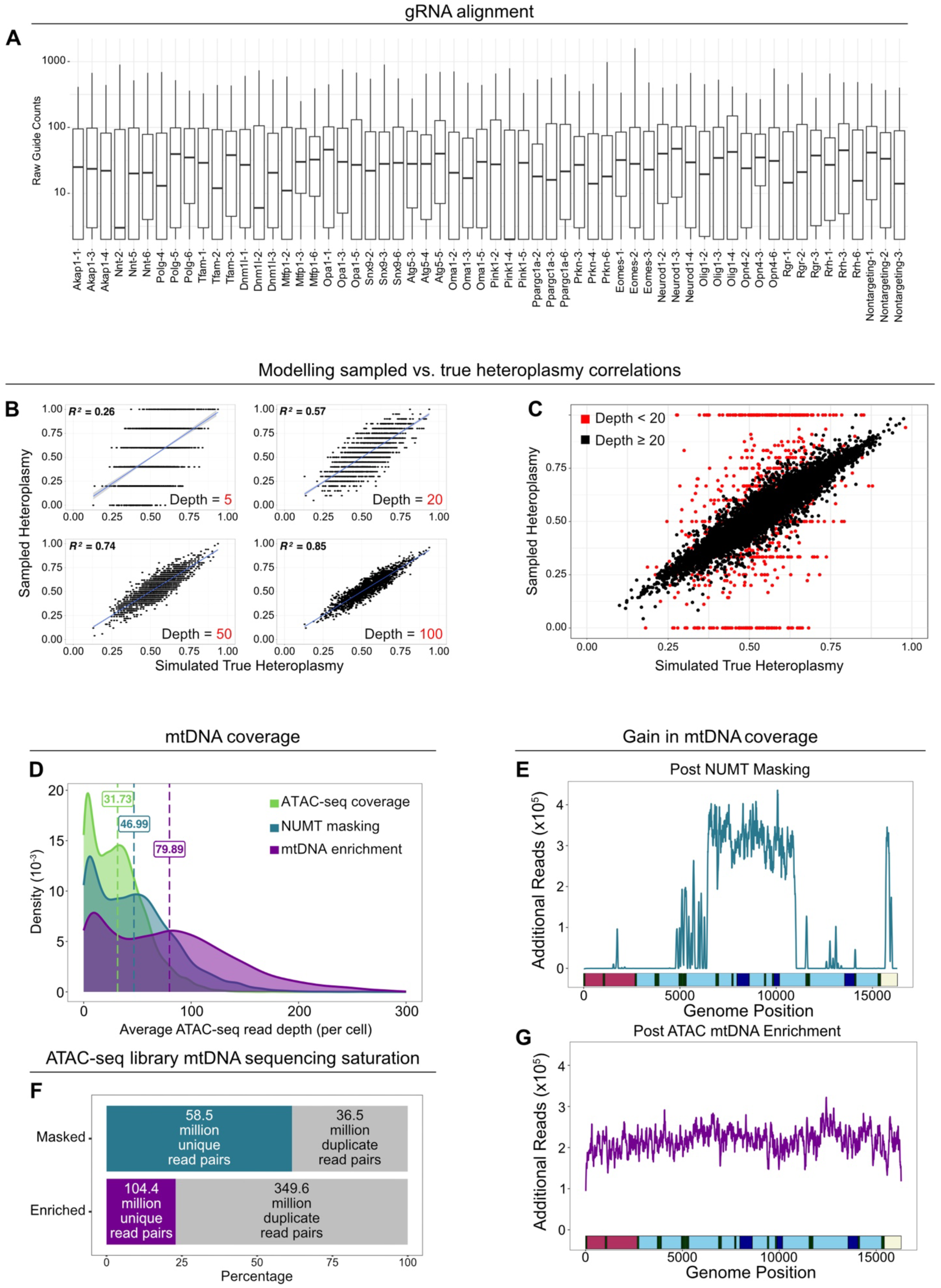
gRNA detection and targeted enrichment of mtDNA. (A) Raw read counts per cell aligned to each of the gRNA sequences included in the gRNA library. For each gRNA, all cells with at least 2 counts are shown. Boxes indicate upper and lower quartiles and mid-line indicates median. (B) Sampled heteroplasmy vs. true heteroplasmy correlation at different mtDNA coverage. A population of 2,000 cells was simulated, and randomly sub-sampled at increasing sequencing depth, with sampled heteroplasmy calls showing better correlation with true heteroplasmy as sample depth increased. Shaded regions around the regression line are 95% CI. (C) Sampled heteroplasmy vs. true heteroplasmy in an *in silico* simulated version of the MitoPerturb-Seq dataset. A population of 6,551 cells was simulated, and sub-sampled using the same mtDNA coverage distribution as obtained from post-mtDNA enrichment mgatk analysis. Cells sampled at depth < 20 are highlighted in red, confirming that a depth threshold of ≥ 20 successfully removed cells with the most inaccurate heteroplasmy calls. (D) Distribution of mtDNA coverage in the MitoPerturb-Seq dataset following alignment of ATAC-seq reads to the standard mm10 reference (light green) and to a NUMT-masked version of mm10 pre- (dark green) and post- (purple) mtDNA hybridization capture-based enrichment. Mean mtDNA coverage values for each approach are indicated. (E) Per-base additional mtDNA coverage gained when aligning ATAC-seq reads to a NUMT-masked version of the mm10 genome compared to the standard mm10 reference. (F) Percentage of unique vs duplicate sequencing reads in pre- and post-mtDNA enrichment ATAC-seq libraries. Raw read-pair numbers for each library are indicated. (G) Line plot showing per-base additional mtDNA coverage gained following hybridization capture enrichment of mtDNA-specific sequences, aligned to the NUMT-masked mm10 genome.

**Figure S3.**
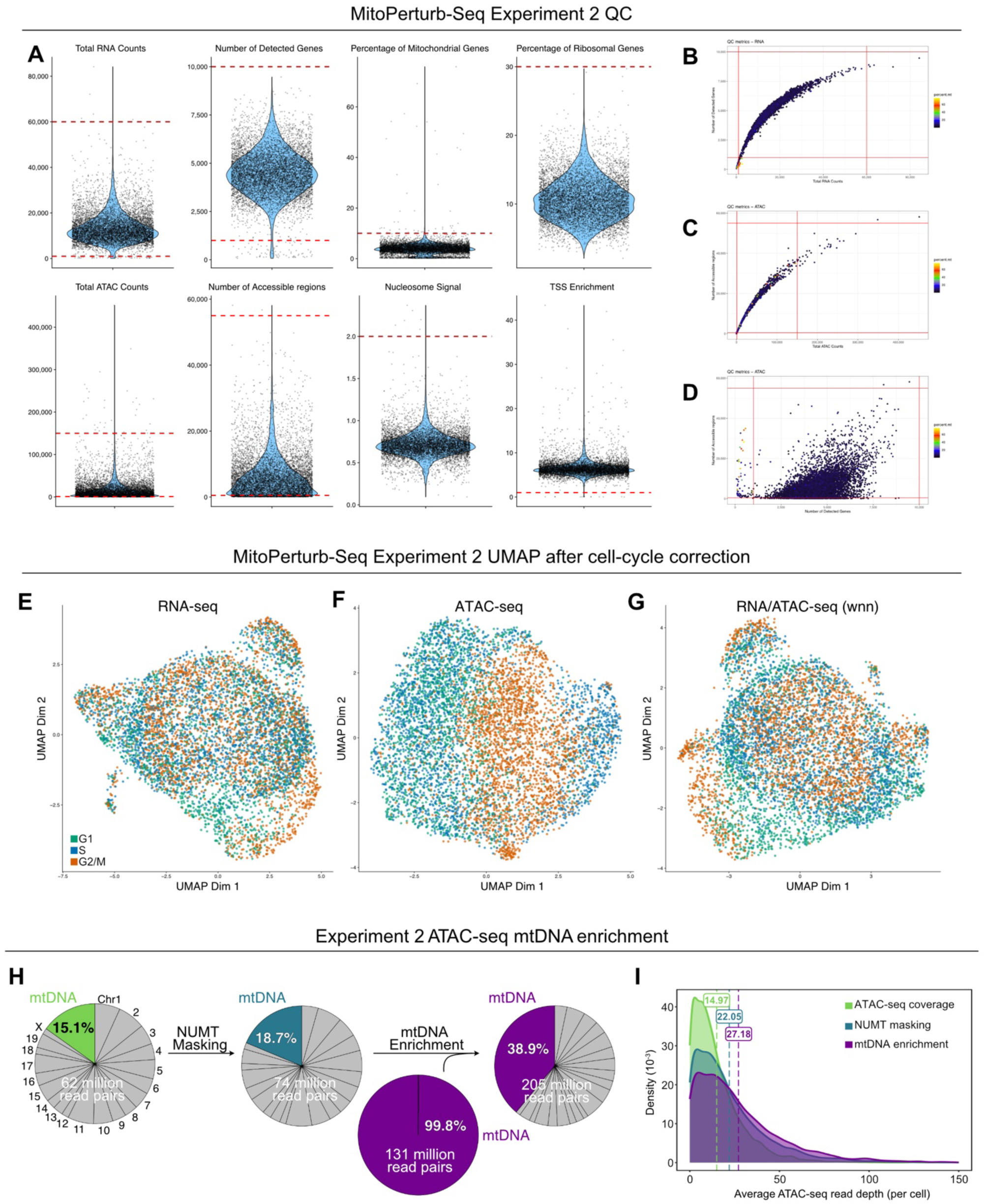
Quality control steps of experiment 2. (A) QC violins from Seurat for RNA-seq and ATAC-seq, dashed red lines indicate thresholds used for filtering, as described in the methods. (B-D) QC plots of nCountRNA (B), nCountATAC (C) and nFeatureRNA (D) versus nFeatureRNA (B) or nFeatureATAC (C,D), colored by percent mitochondrial reads, with lines indicating cutoff thresholds shown in violin plots in (A). (E-G) UMAPs showing cells from the MitoPerturb-Seq dataset in Experiment 2, clustered based on overall RNA expression patterns (E), chromatin accessibility profiles (F) and a combined Weighted Nearest Neighbor (WNN) analysis (G), following regression of cell cycle heterogeneity from the RNA-seq dataset. Each point represents an individual cell. Cells are colored by cell cycle phase. (H,I) Percentage of ATAC-seq reads (H) and distribution of mtDNA coverage (I) in the replicate MitoPerturb-Seq dataset aligning to the mtDNA using the standard mm10 reference (light green), a NUMT-masked version of the mm10 genome (dark green), and following addition of enriched mtDNA reads following hybridization capture (purple). Mean coverage values for each group are indicated in (I).

**Figure S4.**
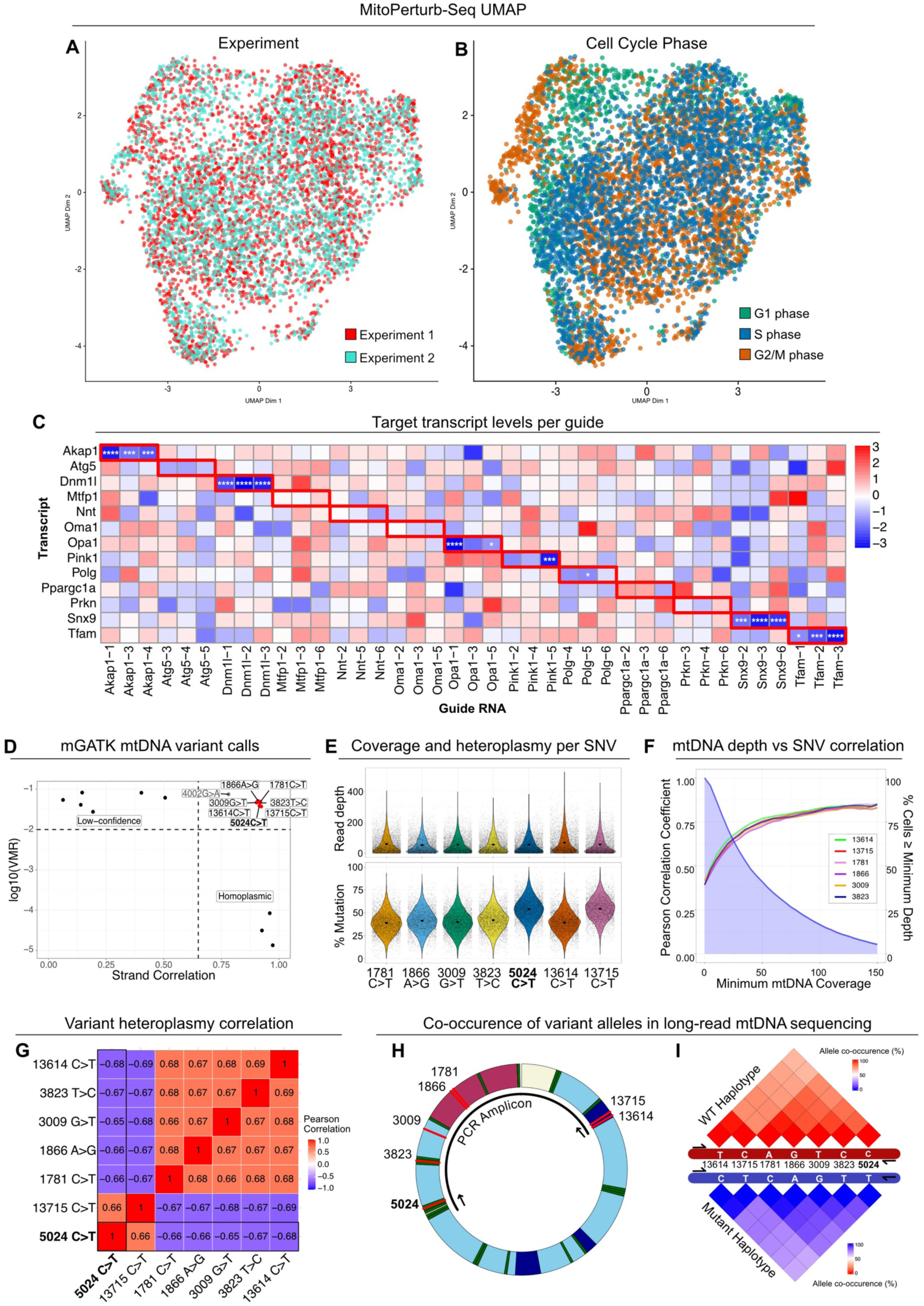
Data-integration, gRNA efficiency, co-segregation of heteroplasmic mtDNA variants. (A,B) UMAPs of the integrated MitoPerturb-Seq dataset, clustered based on WNN analysis following regression of cell cycle heterogeneity from the RNA-seq dataset. Cells are colored by technical replicate (A) or cell cycle phase (B). (C) Scaled transcript levels for each candidate target gene across all perturbation groups, split by individual gRNA assignment. Expression values normalized on a per-column basis. Two-tailed t-tests comparing target transcript expression in the relevant perturbation group vs NT gRNA group, * p < 0.05, *** p < 0.001, **** p < 0.0001. (D) High-confidence heteroplasmic variants present in the m.5024C>T MEF cells, identified by mgatk. Seven variants (red) were previously reported in this mouse strain, with an additional variant, m.4002G>A, that was previously unreported and likely to be a clone-specific *de novo* mutation not present in the mouse strain. (E) Per-cell read depth (top) and heteroplasmy (bottom) at each of the seven main variant sites (SNVs). (F) Pearson correlation coefficients (left axis) between heteroplasmy calls at each of the indicated SNVs and the m.5024C>T site with increasing minimum read depth threshold (horizontal axis). Blue shaded area indicates the percentage of cells above the depth threshold (right axis). Correlation between SNVs increased with more accurate heteroplasmy calls at higher read-depths. (G) Pearson correlation between all variant sites with minimum mtDNA depth threshold >20. (H,I) Schematic of the mtDNA showing the location of the seven main SNVs and the long-range PCR amplicon (H) used for long-read sequencing (I). Percentage of long-reads with cooccurrence of each allele with any of the other alleles on the same haplotype (WT, red, top; mutant, blue, bottom). Co-occurrence is less strong for SNVs further away from each other, due to PCR-mediated recombination during long-range PCR. mtDNA color code: purple = rRNA genes; green = tRNA genes; light/dark blue = protein coding genes; cream = non-coding region; red = SNVs.

**Figure S5.**
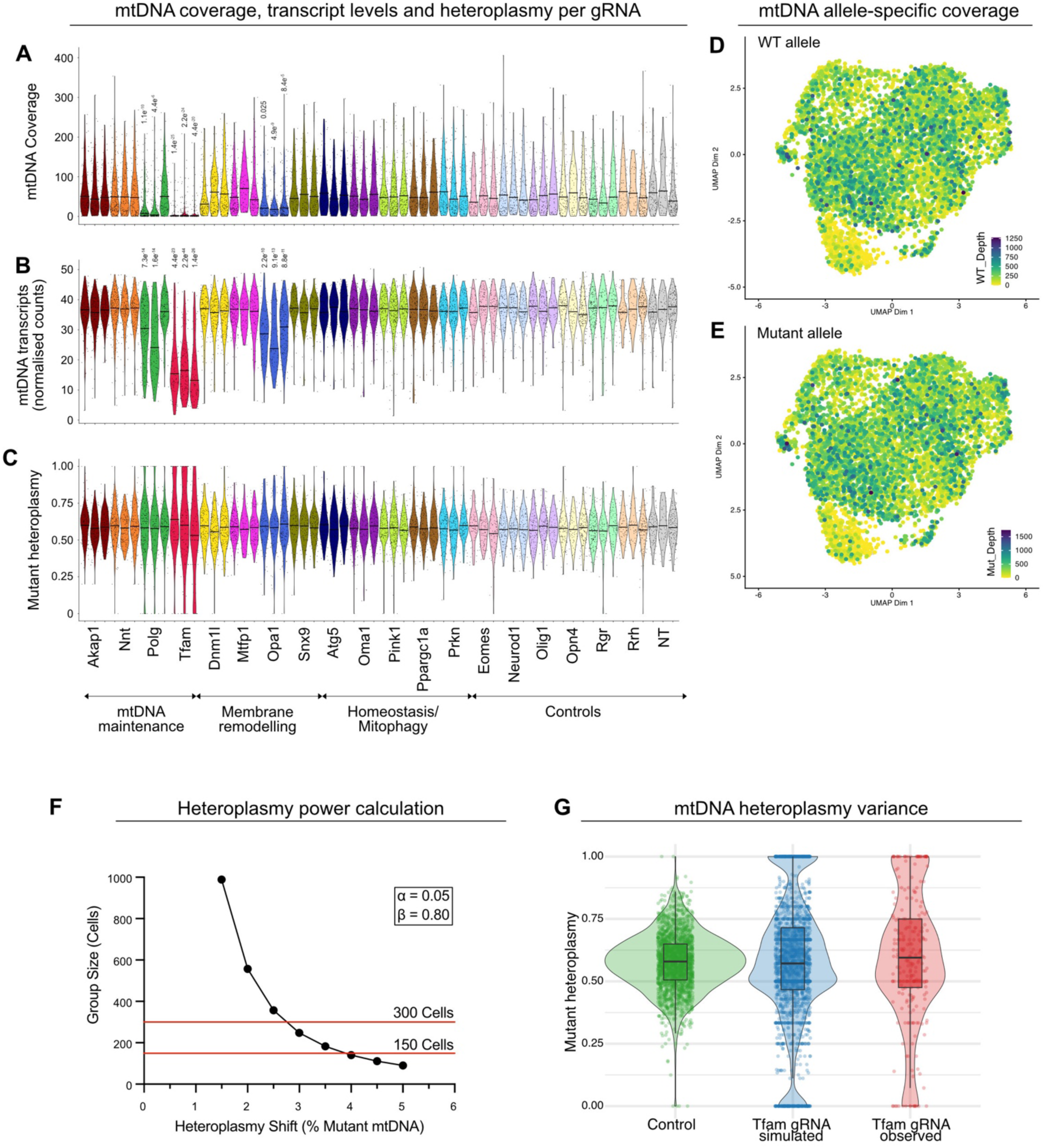
MitoPerturb-seq identifies mtDNA depletion following targeted gene perturbation. (A-C) Per-cell mtDNA coverage (A), mtDNA transcript levels (B) and heteroplasmy (C) for each gRNA group, including controls. Each violin represents the data from one of the three gRNAs targeting the corresponding gene. Pairwise t-tests with multiple testing correction (Bonferroni), p-values specified in figure, all values < 0.05 are shown. Red lines indicate median (A,B) or mean (C). (D,E) UMAPs showing per-cell mtDNA coverage of the WT (D) and mutant (E) allele. (F) Line plot showing the group size required to confidently identify mean heteroplasmy shifts of various magnitudes at a power >0.8 and p<0.05. A mean group size of 179 cells would be sufficient to identify a mean heteroplasmy shift of 3.5 - 4 %, assuming all cells shifted in the same direction. (G) Single-cell mtDNA heteroplasmy levels and distribution in control (green) and Tfam (red) gRNA cells, and in simulated Tfam gRNA cells (blue).

**Figure S6.**
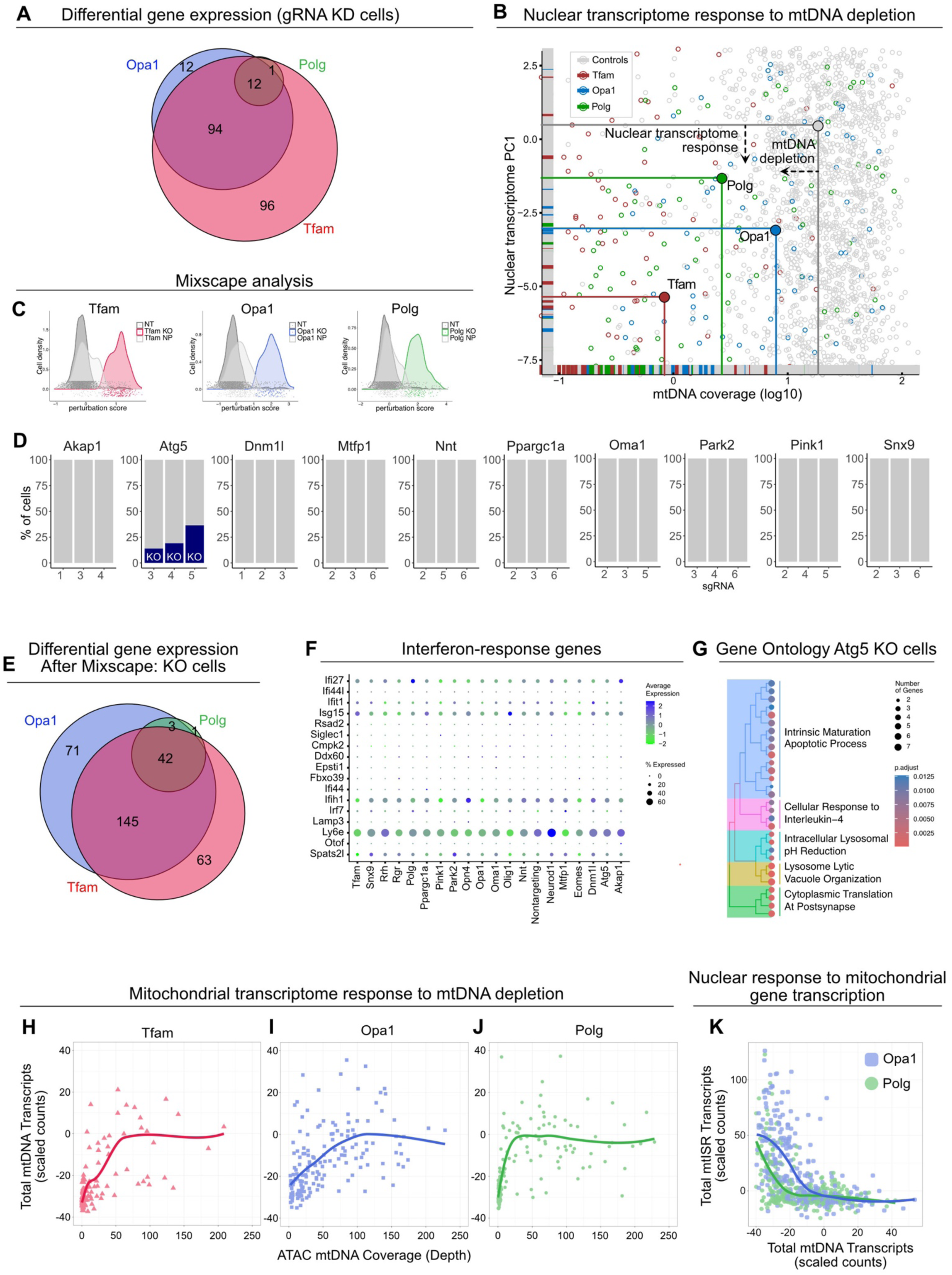
Differential gene expression upon gRNA perturbation. (A) Overlap of DEGs identified in the Tfam, Opa1 and Polg perturbation groups. (B) Strength of principal component (PC1) of nuclear transcriptome in all cells plotted against the mtDNA coverage (log10 transformed), with Polg, Tfam and Opa1 perturbation groups and Controls highlighted, before Mixscape analysis. (C,D) Perturbation scores of cells in the Tfam, Opa1 and Polg gRNA groups (C) and the proportion of cells assigned one of the three gRNAs targeting the indicated gene (D) following Mixscape analysis. Remaining cells (grey) were assigned as non-perturbed (NP). (E) Overlap of DEGs identified in the Tfam, Opa1 and Polg perturbation groups after Mixscape analysis. (F) Average expression of interferon-response genes in cells across the various perturbation groups, indicating overall low expression of these genes, and no activation in any of the groups. (G) Significantly enriched Gene Ontology (GO) terms, grouped by biological process, identified in the Atg5 KO group following Mixscape analysis. Full details of the identified GO terms are in Table S5. (H-J) Per-cell mtDNA transcript levels against mtDNA coverage for the Tfam (H), Opa1 (I) and Polg (J) perturbation groups. (K) mtDNA transcript levels against combined transcript levels of genes involved in the mitochondrial integrated stress response in the Opa1 and Polg perturbation groups, indicating increased activation of the stress response in Opa1 KO cells compared to Polg KO cells at similar mtDNA transcript levels.

**Figure S7.**
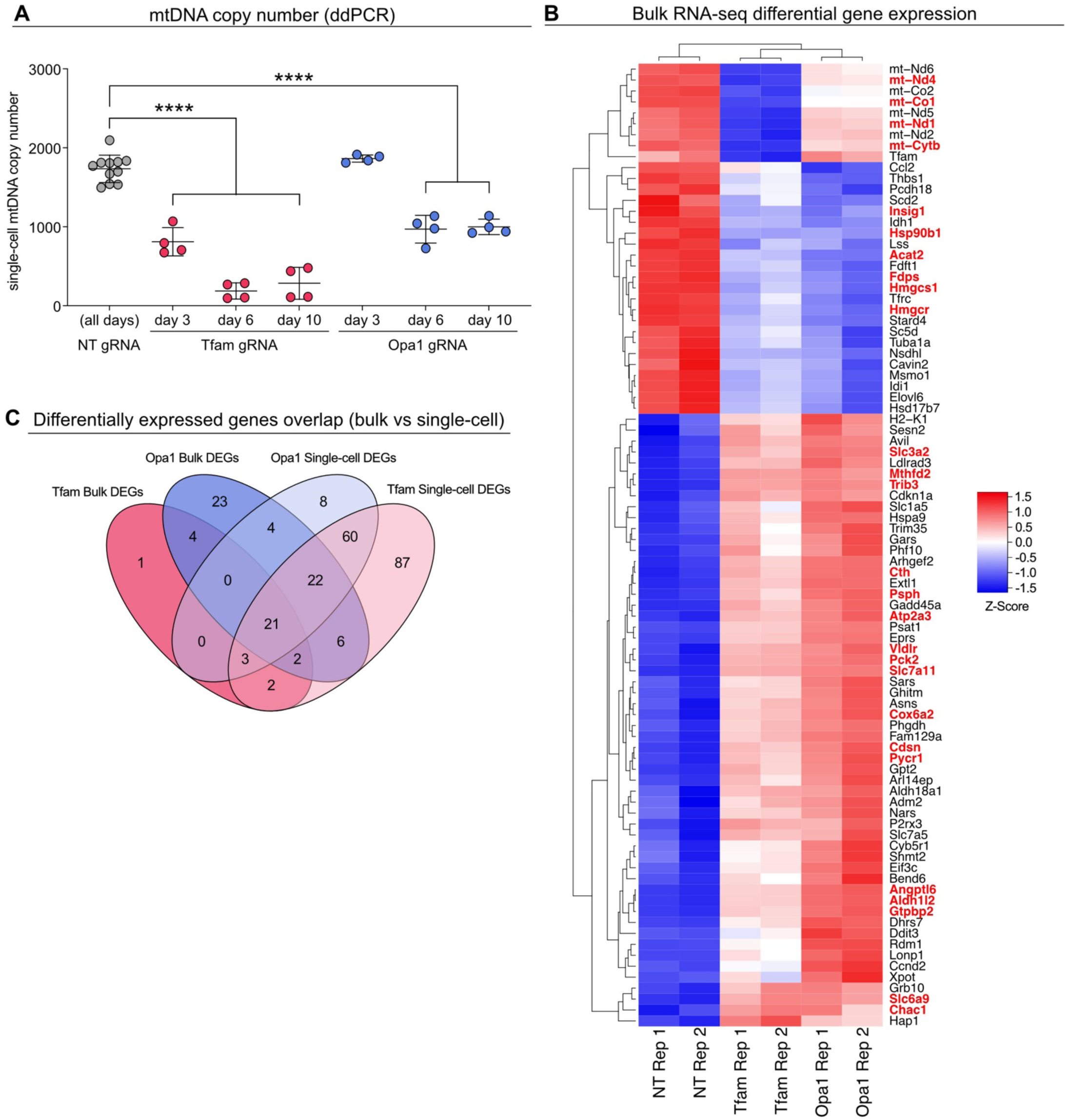
Bulk RNA-seq upon targeted perturbation of Tfam and Opa1. (A) Timecourse of mtDNA CN in m.5024C>T MEFs at 3 days, 6 days and 10 days following transduction with non-targeting (NT, grey), Tfam (red) or Opa1 (blue) gRNAs. Each point represents a 20-cell bulk ddPCR measurement normalized to cell number. One-way ANOVA with Tukey’s post-hoc test, pairwise comparisons to NT control, **** p < 0.0001. (B) Relative expression of DEGs in bulk RNA-seq data from Tfam- or Opa1-gRNA cells compared to non-targeting controls. All genes shown were significantly differentially expressed (FDR < 0.05 and log2-fold-change > 0.25) in at least one gRNA condition; genes highlighted in red were differentially expressed in both Tfam- and Opa1-gRNA cells. (C) Overlap of DEGs identified in bulk RNA-seq and MitoPerturb-Seq Tfam and Opa1 KD cells.

**Figure S8.**
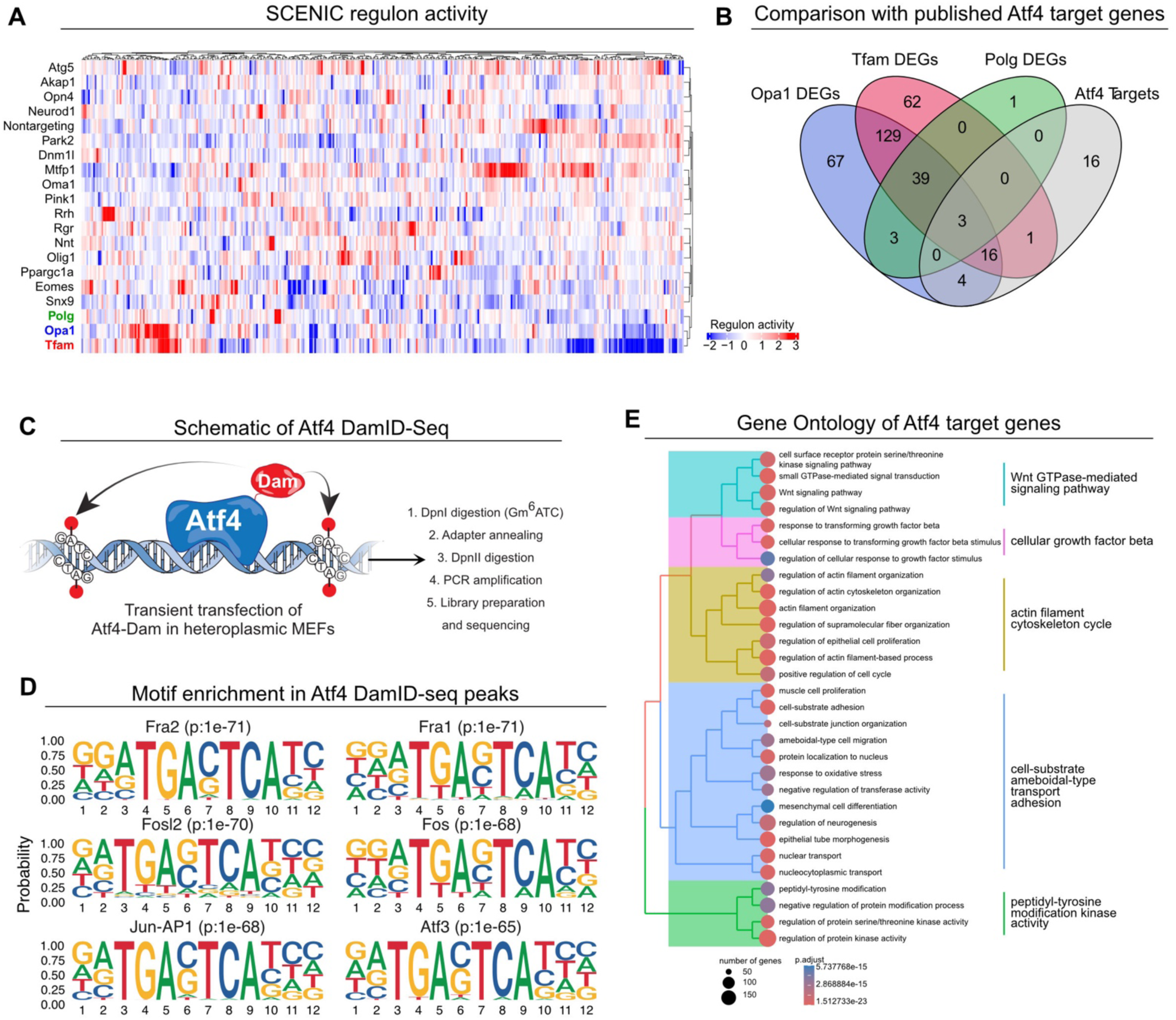
Atf4 DamID-seq in heteroplasmic MEFs. (A) Regulon activity following SCENIC analysis. Average AUC scores were calculated for each gRNA group; all regulons identified are shown, and also listed in Table S6. (B) Intersection of Tfam, Opa1 and Polg KO MitoPerturb-Seq DEGs (post-Mixscape analysis) with a published list of high-confidence Atf4 target genes. (C) Schematic of Atf4-DamID-seq in heteroplasmic MEFs. (D) *De novo* motif enrichment using HOMER in Atf4-bound regions. (E) Significantly enriched Gene Ontology (GO) terms, grouped by biological process, identified in the Atf4 target genes.

**Figure S9.**
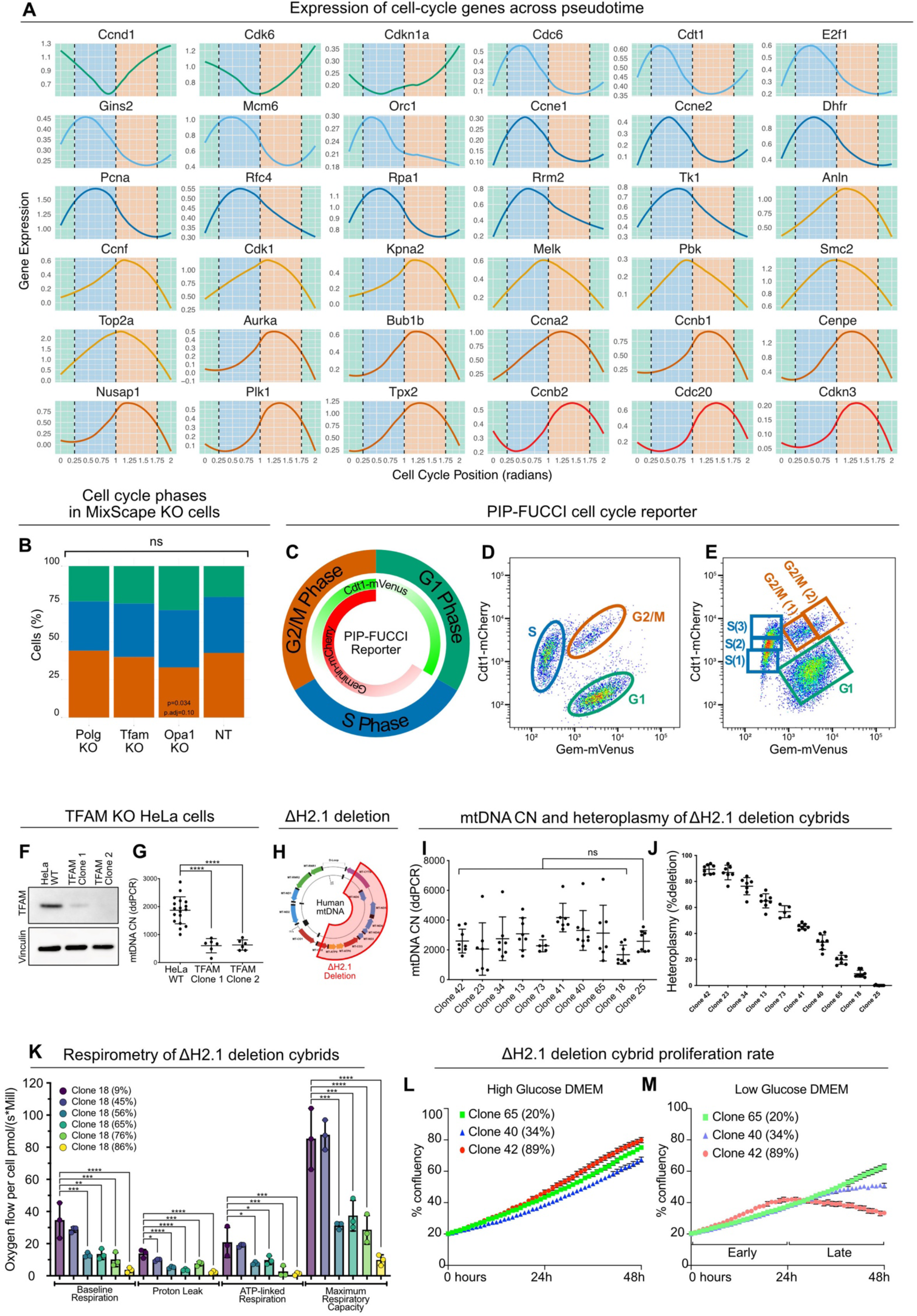
Cell cycle slowing in response to mtDNA depletion. (A) Expression profiles of indicated cell-cycle genes across cell-cycle pseudotime determined by tricycle. (B) Percentage of cells in each cell-cycle phase in Tfam, Opa1 and Polg KO cells, post-Mixscape analysis, compared to non-targeting gRNA cells. Chi-squared tests with multiple testing correction (Bonferroni); ns, not significant. (C) Schematic showing the expression patterns of the PIP-FUCCI fluorescent reporters Cdt1-mVenus and Geminin-mCherry across the cell-cycle. (D,E) Flow cytometry plots showing the gating strategies used to isolate cells for mtDNA CN measurements based on cell-cycle phase for both inter-phase (D) and intra-phase (E) comparisons. (F,G) Western blot (F) and ddPCR measurements (G) showing KD of TFAM at the protein level (F) and reduced mtDNA CN (G) in HeLa cells transduced with two independent TFAM gRNAs. Vinculin is shown as a loading control (F). One-way ANOVA with Tukey’s post-hoc test, **** p < 0.0001. (H) Map of the human mtDNA, highlighting the 7.5 Kb deletion present in the DeltaH2.1 cybrid cell line. (I,J) mtDNA CN (I) and heteroplasmy (J) of eight clonal populations isolated from the DeltaH2.1 cybrid cell line. One-way ANOVA of the mtDNA CN distributions was significant (p = 0.015); pairwise comparisons of all heteroplasmic clones to Clone 25 (0% heteroplasmy) using Tukey’s post-hoc test, showed no significant differences in mtDNA CN; ns, not significant. (K) Respirometry of DeltaH2.1 cybrid cells with increasing heteroplasmy levels, using Oroboros, showing baseline respiration (BR), proton-leak (PL), ATP-linked respiration (AR) and maximal respiratory capacity (MC). One-way ANOVA with Tukey’s post-hoc test performed for each tested parameter, all significant pairwise comparisons are shown. * p < 0.05, ** p < 0.01, *** p < 0.001, **** p < 0.0001. (L,M) Growth curves of DeltaH2.1 cybrid clones carrying different levels of heteroplasmy in high (L) and low (M) glucose culture medium.

**Figure S10.**
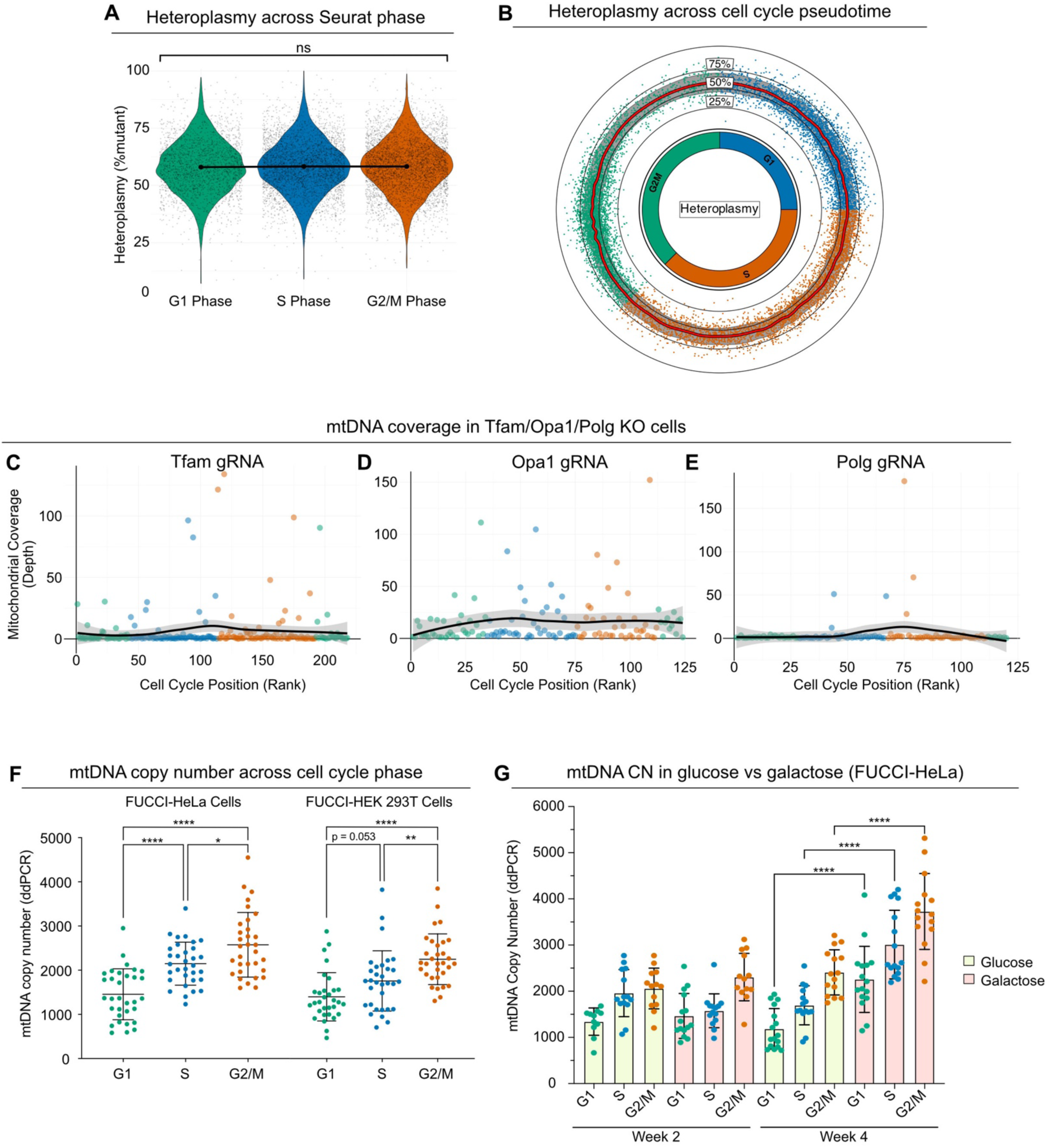
Relaxed replication of mtDNA. (A,B) Heteroplasmy levels across cell cycle stages using Seurat (A) or across cell cycle pseudotime based on tricycle pi score (B). Red line = mean, grey band = standard deviation. One-way ANOVA with Tukey’s post-hoc test; ns, not significant. (C-E) mtDNA coverage across cell cycle pseudotime with cells ranked based on tricycle pi score for the Tfam (C), Opa1 (D) and Polg (E) perturbation groups. (F) Single-cell mtDNA CN measurements in PIP-FUCCI-expressing HeLa and HEK293T cells following flow sorting based on cell cycle phase. Error bars are mean ± SD. One-way ANOVA with Tukey’s post-hoc test performed across cell-cycle phases for each cell type, * p < 0.05, ** p < 0.01, **** p <0.0001. (G) Single-cell mtDNA CN measurements in PIP-FUCCI-expressing HeLa cells following flow sorting based on cell cycle phase. Cells were cultured in DMEM supplemented with either high-glucose or galactose for four weeks. Error bars are mean ± SD, one-way ANOVA with Tukey’s post-hoc test performed per cell-cycle phase across time points and culture conditions, **** p <0.0001.

**Supplementary Video 1:** PIP-FUCCI-expressing HeLa cells time-lapse movie at 20-minute intervals over 24 hours. Related to **Figure 4**.

## Supplementary tables

**Supplementary table 1:** gRNA sequences used in this study, including gRNAs used for the single gRNA transductions and bulk RNA-sequencing.

**Supplementary table 2:** Hybridization capture probes for enrichment of mouse mtDNA from ATAC-seq libraries.

**Supplementary table 3:** Differentially expressed genes (log2FC > 0.25, p.adj < 0.05) from all perturbation groups.

**Supplementary table 4:** Differentially expressed genes (log2FC > 0.25, p.adj < 0.05) in Tfam, Opa1, Polg and Atg5 KO cells post-Mixscape analysis.

**Supplementary table 5:** Gene ontology for differentially expressed genes in Tfam, Opa1, Polg and Atg5 KO cells.

**Supplementary table 6:** Transcription factor regulon activity scores (scaled), identified by SCENIC analysis of all perturbation groups.

**Supplementary table 7:** Atf4 DamID-seq consensus peak coordinates.

**Supplementary table 8:** Atf4 DamID-seq bound genes in heteroplasmic MEFs.

## Notes

### Competing Interest Statement

The authors have declared no competing interest.

